# Deconstructing the individual steps of vertebrate translation initiation

**DOI:** 10.1101/811810

**Authors:** Adam Giess, Yamila N. Torres Cleuren, Håkon Tjeldnes, Maximilian Krause, Teshome Tilahun Bizuayehu, Senna Hiensch, Aniekan Okon, Carston R. Wagner, Eivind Valen

## Abstract

Translation initiation is often attributed as the rate determining step of eukaryotic protein synthesis and key to gene expression control ^1^. Despite this centrality the series of steps involved in this process are poorly understood ^2,3^. Here we capture the transcriptome-wide occupancy of ribosomes across all stages of translation initiation, enabling us to characterize the transcriptome-wide dynamics of ribosome recruitment to mRNAs, scanning across 5’ UTRs and stop codon recognition, in a higher eukaryote. We provide mechanistic evidence for ribosomes attaching to the mRNA by threading the mRNA through the small subunit. Moreover, we identify features regulating the recruitment and processivity of scanning ribosomes, redefine optimal initiation contexts and demonstrate endoplasmic reticulum specific regulation of initiation. Our approach enables deconvoluting translation initiation into separate stages and identifying the regulators at each step.

## Introduction

In eukaryotes, translation initiation is a highly orchestrated sequence of events where the ribosomal 43S pre-initiation complex (PIC) is first recruited to the beginning of the transcript through interactions with initiation factors and the 5’ m^7^G cap ^4^. The 43S PIC then scans the transcript in a 5’-to-3’ direction until a suitable translation initiation site (TIS) is encountered. Upon recognition of the TIS, the large ribosomal subunit is recruited to form an elongation-capable 80S ribosome. These initiation steps are broadly acknowledged to be a rate limiting factor in protein synthesis ^3,5,6^. Despite this, our knowledge of ribosome recruitment, scanning, and TIS recognition is limited.

Ribosome profiling (ribo-seq) has enabled global quantification and localization of translation through the capture of footprints from elongating 80S ribosomes ^7^. A limitation of ribo-seq, however, is that it is blind to ribosomes from other stages of translation. Recently, translation complex profiling (TCP-seq) was introduced which circumvented this problem by crosslinking all stages of ribosomes to the mRNAs ^8^. However, because this technique relies on first purifying 80S ribosome containing transcripts, it is limited to studying 40S-ribosome positioning to transcripts that have at least one 80S ribosome and are thus actively translated. Here, we have expanded this approach to capture footprints from all ribosome-associated mRNAs, including transcripts not bound by any 80S subunit. Our approach immobilizes all ribosomal subunits on the mRNA by paraformaldehyde crosslinking, followed by sucrose gradient separation of the small subunits from the 80S complexes ^8^ (Fig. 1A). After extracting the RNA, sequencing libraries are made of each fraction using template-switching, which enables the use of ultra-low input material (1 ng) ^9^. Because our method captures different populations of ribosomes than TCP-seq, we will refer to our modified protocol as “Ribosome Complex Profiling” (RCP-seq).

**Figure 1:**
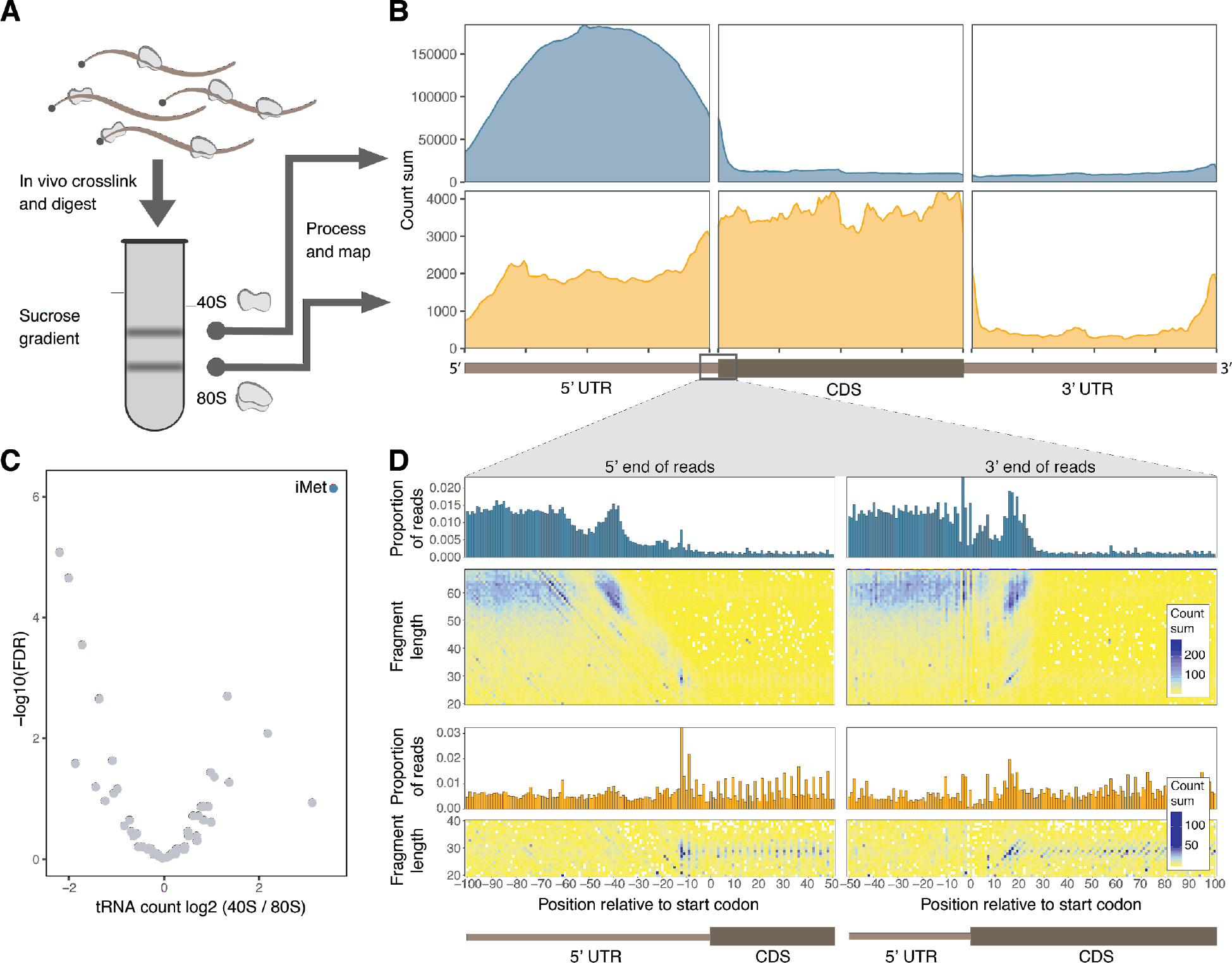
RCP-seq selectively captures 80S ribosomes and small subunits in zebrafish. **A**) Schematic representation of RCP-seq protocol. **B**) Coverage of RCP-seq reads across all transcripts. Footprints from small subunits (blue) map predominantly to 5’ UTRs while 80S footprints (orange) map predominantly to coding regions (CDS). **C**) Abundance of tRNA species (x-axis) and false discovery rate (FDR) (y-axis) between the RCP-seq small subunit (40S) and 80S fractions. Initiator Met-tRNA is highlighted (blue). **D**) Over-representation of RCP-seq small subunit (upper) and 80S (lower) fraction footprints around start codons. Counts of 5’ (left) or 3’ (right) ends of fragments are summed across highly expressed genes (>= 10 FPKM). The barplots show the proportion of read counts per position (x-axis), while the heatmaps show the same counts stratified by length (y-axis) and coloured by total count.

Here, we use RCP-seq to capture footprints of both 80S ribosomes and small ribosomal subunits across the transcriptome of a developing zebrafish embryo (Fig. 1A, Methods). Mapping scanning small subunits over 5’ UTRs allows us to distinguish three distinct phases during translational initiation: 1) recruitment of small subunits to the mRNAs, 2) progression to the start codon, and 3) conversion of scanning to elongating ribosomes.

## Results

To investigate the regulation of translation initiation in a vertebrate, we performed RCP-seq during zebrafish embryo development (see Methods and Supplementary Notes). As expected under the scanning model of translation, the footprints from the small subunit fraction predominantly mapped to the 5’ UTR of the transcripts, while the elongating 80S footprints mapped to the protein coding region (CDS). A sharp divide between the fractions occurred at the start codon consistent with the conversion of scanning 43S PICs to elongating 80S ribosomes (Fig. 1B). Here, the distribution of footprint lengths also revealed a range of ribosomal initiation conformations similar to those previously reported in yeast (Fig. 1D, Fig. S1) ^8^. As previously reported for TCP-seq, tRNA species contained within ribosomes are also selectively protected by RCP-seq, and consistent with capturing scanning ribosomes we found initiator MET-tRNA strongly enriched in the small subunit fraction (Fig. 1C). Taken together, these observations provide strong support for the selective capture of footprints from small subunits with RCP-seq.

We first sought to understand how the 43S PIC is recruited to the mRNA. Previous studies have suggested two alternative models for the 43S PIC binding to mRNA ^2^. In the first, mRNA is “threaded” through the mRNA channel of the complex while in the second, mRNA “slots” directly into the channel, possibly leading to suboptimal scanning of the first nucleotides (Fig. 2A) ^2^. The slotting and threading models are predicted to lead to substantially different profiles of protected fragments over the 5’ end (Fig. 2A). Zebrafish mRNAs have a strong enrichment of small subunit footprints coinciding with the 5’ end of transcripts (Fig. S2). The 5’ peak is not present in non-coding RNAs arguing that it is a feature only of translated RNA molecules and not an artifact of the method (Fig. S3). The start of these footprints all coincide with the transcription start site, and have a wide range of read lengths from the lower detection limit (~15 nt) up to about 80nt which is slightly longer than scanning 43S PICs (Fig. 2B and Fig. S2, S4). The majority of footprints downstream of this peak corresponds to the range commonly reported for 43S PICs (60-70 nt) ^8,10^. A similar pattern was also observed when realigning data from TCP-seq in yeast to high-resolution mapping of transcription start sites ^11^ (Fig. S5, Methods). These patterns of increasing lengths of small subunit footprints at the 5’ end of the transcript up to the size of the longest small subunit footprints are consistent with footprints from successive threading of the transcript through the mRNA channel of the 43S PIC complex.

**Figure 2:**
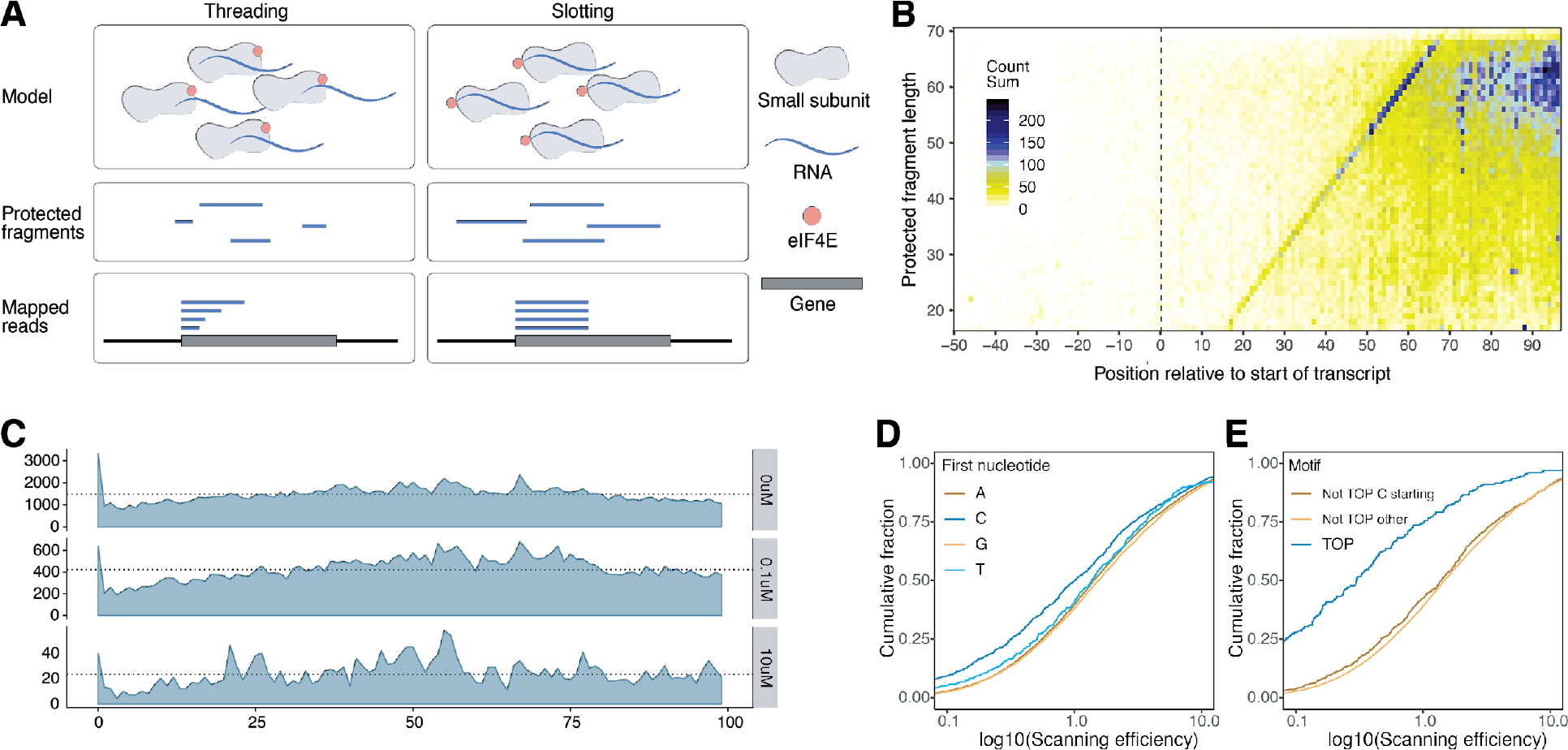
43S PIC recruitment and impact of 5’ transcript features. **A**) Schematic representation of two canonical recruitment models (upper panel): “Threading” (left) and “Slotting” (right), the resulting protected fragments (middle panel) and the location of the mapped reads relative to the transcription start site (bottom panel). **B**) Heatmap of counts from 3’ ends of small subunit reads stratified by length (y-axis) over each position (x-axis) relative to transcription start site. Dotted line shows the beginning of the transcript with all reads ending at the enriched diagonal start. **C**) Small subunit 5’ counts, relative to transcription start site. Dotted lines show the median values per condition (3 conditions shown, control, 0.1 μM and 10 μM 4Ei-10 treatment). **D-E**) Empirical cumulative density of scanning efficiency for highly expressed transcripts (>10 FPKM). **D**) Transcripts colored by their first nucleotide. An initial pyrimidine (C/T) results in lower scanning efficiency than a purine (A/G) (C: p < 2.2×10^−16^, T: p < 2.2×10^−16^; number of transcripts per group, A=10388, C=1663, G=8335, T=967). **E**) Transcripts starting with a TOP motif show reduced scanning efficiency (p < 2.2×10^−16^), while non-TOP transcripts starting with a C show reduced SE but small effect compared to the other non-TOP transcripts (number of transcripts per group Not TOP C starting = 1257, Not TOP other = 19690, TOP = 406).

Under the threading model, the cap-binding initiation factor eIF4E is placed at the leading edge of the 43S PIC and mRNA is threaded through the mRNA-binding channel ^2^. To test the response of the 5’ end peaks to eIF4E inhibition, we sequestered eIF4E using the small molecule inhibitor 4Ei-10 ^12^ (a cell-permeable prodrug improving upon 4Ei-1 ^13^) thereby specifically blocking eIF4E-cap binding, leading to a small but general inhibition of translation (Fig. S6, Supplementary Notes) followed by RCP-seq. This resulted in a global depletion of 5’ peaks with the ratio of reads at the 5’ peak relative to reads internal to the 5’ UTR reduced to ~63% of WT levels. (Fig. 2C, Fig. S3, S7). In transcripts with very short 5’ UTRs only threading is expected to be able to initiate translation as slotting would deposit the small subunit too far downstream to scan the start codon ^2,14^. Consistently, we observed a strong 5’ peak reduction in response to eIF4E inhibition in transcripts initiated through the Translation Initiator of Short 5’ UTR (TISU) motif (Fig. S3E-F), in line with previous reports that these transcripts are eIF4E-sensitive ^14^. Collectively, this suggests threading is dependent on eIF4E and is a common recruitment pathway during early development.

We next asked which features could influence the recruitment of 43S PICs to the 5’ cap. To measure the amount of 43S PICs present on 5’ UTRs, we defined the scanning efficiency (SE) as the number of small subunit footprints over a 5’ UTR relative to its mRNA abundance (Fig. S8). This metric is conceptually identical to the widely used translational efficiency (TE), which measures elongating ribosomes relative to mRNA abundance ^15^ (Supplementary Notes). Using this metric, we observed that transcripts with a 5’ C, and to a lesser extent 5’ T, showed reduced SE and TE compared to transcripts beginning with an A or G (Fig. 2D, Fig. S9A). This is consistent with biochemical studies which have shown that transcripts beginning with a pyrimidine (C/T) have a lower affinity for eIF4E binding than those starting with a purine (A/G) ^16,17^. An initial C is a feature of transcripts containing a 5’ Terminal Oligo-Pyrimidine (TOP) tract, a motif often present in mRNAs encoding the protein synthesis machinery and a target of mTOR-mediated translation control ^16,17^. We found that while C has an effect, the overall effect on SE and TE reduction is dominated by mRNAs with the TOP-motif (Fig. 2E, Fig. S9B). This demonstrates that during early development a reduced number of 43S PIC are recruited to TOP-motif containing transcripts resulting in reduced translation.

As the 43S PIC progresses through the 5’ UTR, it can encounter obstacles that can lead to termination of scanning. The RCP-seq data revealed this as a slight decline of scanning ribosomes throughout the 5’ UTR (Fig. S10A). Under the assumption that averaged across all transcripts scanning proceeds at a uniform pace throughout the 5’ UTR, we compared the density of small subunit complexes of all transcripts at the 5’ end of the mRNA to the density proximal to the start codon (Fig. S10B). Based on this analysis, we estimate that on average across all transcripts about 68 % of all ribosomes recruited to the 5’ end reach the start codon. The loss of scanning ribosomes is largely contingent on whether the 5’ UTR contains one or more upstream open reading frames (uORF)^18,19^ (Table S1) with only a modest correlation (Spearman’s rho: - 0.03) with 5’ UTR length if you control for number of uORFs (Fig. S11). In transcripts that lack a uORF, scanning overall maintains high processivity, consistent with previous reports ^20^, with a median of 95% of ribosomes retained. Collectively, this argues that scanning is highly stable, and globally regulated through 5’ UTR elements promoting disassociation.

In transcripts containing uORFs, the CDS is translated either from ribosomes that fail to recognise the often sub-optimal uORF TIS ^7,21,22,23^, or by reinitiating ribosomes that continue scanning after translating the uORF ^24,25^. Therefore, uORFs typically lead to reduced protein synthesis by consuming scanning 43S PICs (Fig. 3A-C) ^26^. Consistent with our global estimates, we find a local decline of 43S PIC footprints coinciding with an increase in 80S footprints at uORF TIS’ (Fig. 3D). The ratio of the 43S PIC density upstream vs downstream of a uORF TIS can therefore quantify to what extent uORFs consume scanning 43S PICs (Fig. 3D). As expected, uORFs starting with an ATG start codon (Fig. 3E-F) and with a TIS context similar to the Kozak sequence (Fig. 3E) have the highest 43S PIC consumption.

**Figure 3:**
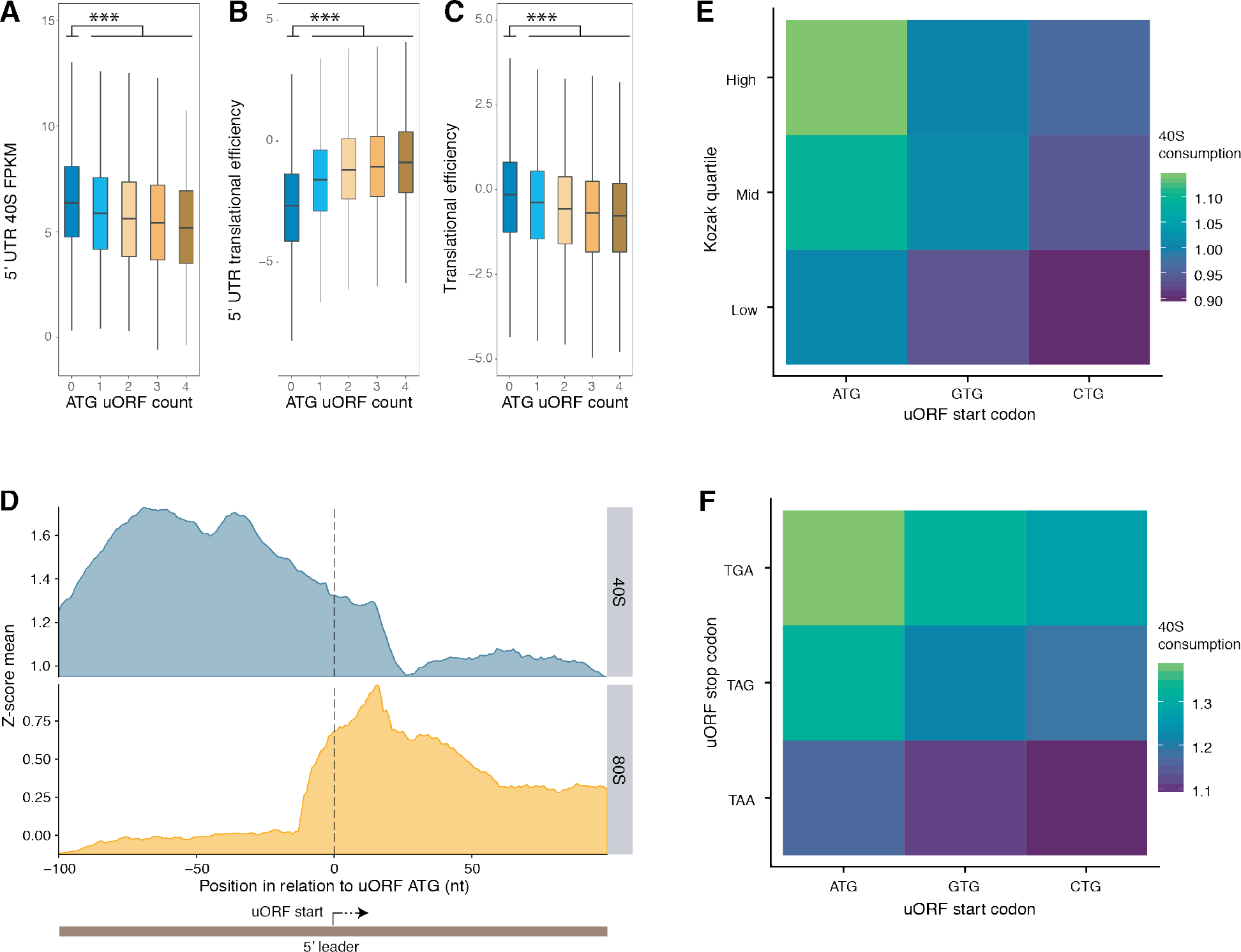
uORFs reduce the number of 43S PICs scanning across 5’ UTRs. **A-C)** The impact of the number of uORFs on (**A**) scanning subunits on 5’ UTR, (**B**) the translational efficiency of the 5’ UTR, and (**C**) the translational efficiency of the protein (*** = p-values < 0.001). **D)** Coverage of small subunit (40S) footprints (upper, blue) and ribo-seq 80S complex footprints (lower, orange) in fixed windows of 100 nt up- and downstream of the first ATG uORF. **E)** Heatmaps showing the rate of scanning subunit consumption as measured by the ratio of small subunit reads upstream versus downstream of all uORFs stratified by surrounding Kozak score and start codon. **F**) Same as E, but with ranking of start and stop codon.

Surprisingly, we found the ability of the small subunit to resume scanning after uORF translation to be highly dependent on the choice of stop codon. For proteins, TAA and TGA have been reported as the most and least efficient termination codons, respectively ^27^. Consistently, uORFs with TGA have the greatest reduction of downstream scanning small subunits (Fig. 3F), the highest density of downstream 80S footprints (Fig. S12A) and the lowest ratio of small subunit to 80S complexes directly over their stop codons (Fig. S12B). This less efficient stop codon recognition globally leads to a small, but significant effect on the TE of the downstream CDS (Fig. S12C). This suggests that failure to recognise a stop codon results in extended uORF translation and decreased rates of reinitiation after the translation of the extended uORF ^28–31^. The choice of uORF stop codon can therefore regulate synthesis of the downstream protein.

Whether a 43S PIC will recognize the TIS and trigger initiation of translation depends on the sequence surrounding the start codon. For many species, studies have defined an optimal consensus sequence for translation initiation (the Kozak sequence ^32,33^), often defined from indirect measures such as sequence conservation ^34^ or reporter protein expression ^35^. Uniquely, RCP-seq enables us to directly measure the average initiation rate (IR) on individual transcripts as the ratio of 80S ribosomes in the CDS to small subunit complexes in the 5’ UTR (Fig. S8). By calculating the median IR of all transcripts containing a specific nucleotide at a specific position, this model revealed that the consensus that maximized IR is identical to the known zebrafish Kozak sequence ^34^ (Fig. 4A). This model, however, considers positions independently and therefore only reflects an average over sequences with high IR and not the efficiency of any particular sequence. To obtain this, we grouped all genes with identical sequence context and ranked these sequences by their median IR (Fig. 4B, S13). The resulting ranking was consistent with a previous assessment of a small number of sequences in zebrafish ^34^, but surprisingly revealed that the Kozak sequence is not the optimal context. The highest scoring sequence was AGGC<u>ATG</u> which differs by two bases (G at −3 and −2). More surprisingly, several sequences that differ strongly to the reported Kozak sequence rank above it. To test this ranking, we constructed GFP mRNA reporters with three different initiation sequences, but otherwise identical: 1) AAGC: a sequence highly similar to the Kozak but with low IR, 2) AAAC: the Kozak sequence previously defined for zebrafish, and 3) TGGA: a sequence differing at 4 bases from the Kozak, but with greater IR. The translational efficiency (see Methods) of these reporters when injected into zebrafish embryos confirmed our direct measurements from IR (Fig. 4C). Taken together with previous reports of weaker than expected correlations between Kozak sequences and translation ^36,37^, this demonstrates that a Kozak-similarity measure does not capture the complexity of start codon recognition, but can be obtained by transcriptome-wide quantification through methods such as RCP-seq.

**Figure 4:**
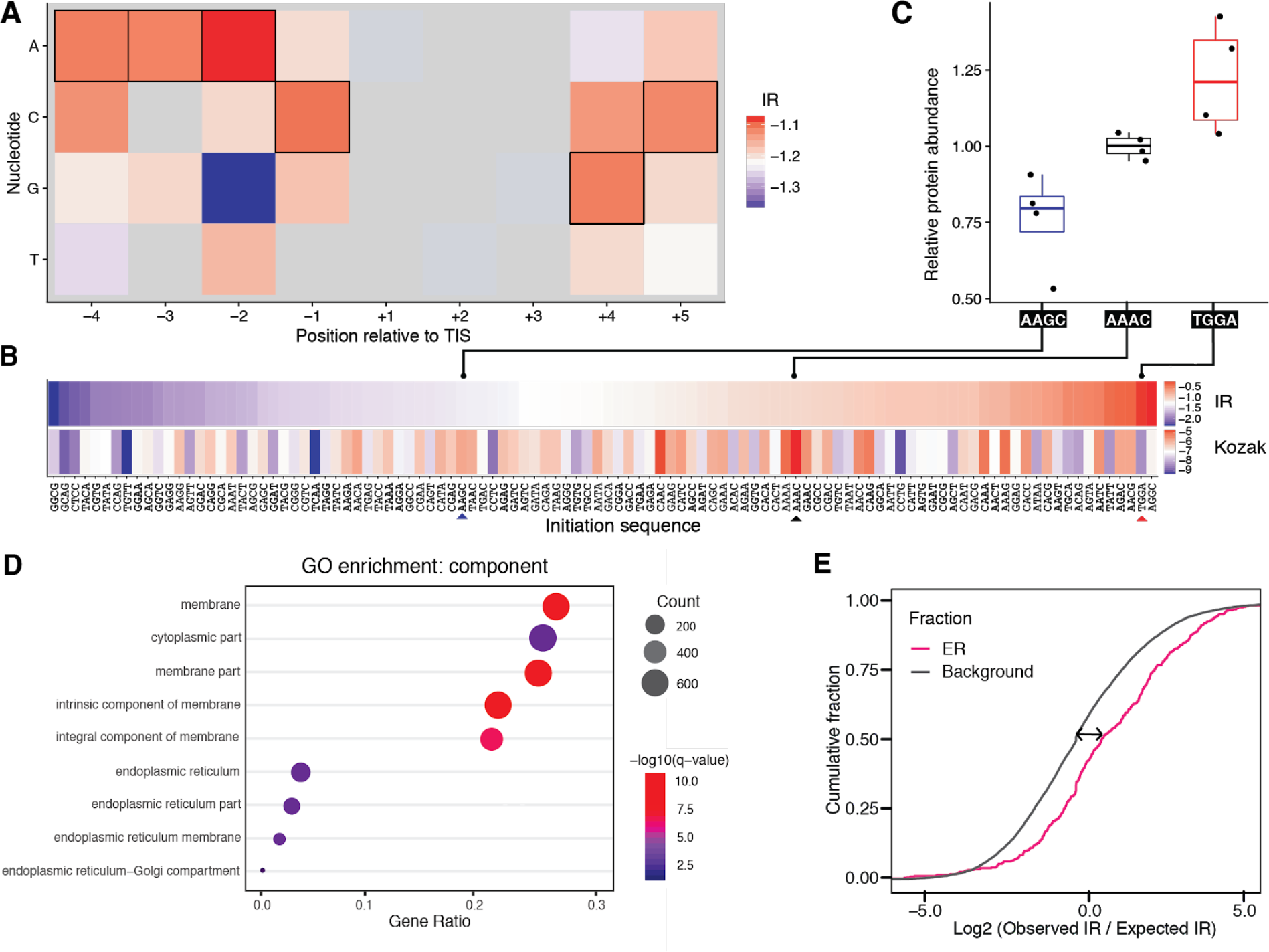
Direct measurements of initiation rate. **A**) Median initiation rates (IR) for all transcripts containing nucleotide (y-axis) at a specific position (x-axis) relative to the protein TIS. The zebrafish Kozak sequence is highlighted with black borders (AAAC<u>ATG</u>GC). **B**) Median IR for entire sequence from −4 to −1 (upper) with corresponding Kozak strength (lower). Arrows indicate the sequences selected for reporter constructs. **C**) Relative protein abundance for GFP reporter constructs for three different initiation contexts, as measured by the protein/RNA abundance ratios in zebrafish embryos at 24 hours post fertilization. **D)** Gene set enrichment analysis of ranked genes based on Initiation Rate (IR) of start contexts. Q-value (color) is based on False Discovery Rate. Gene ratio (x-axis) and size of dots represent the number of genes in each category out of the 2489 included in the analysis. Only significant terms shown. **E)** Cumulative fraction of the fold change of observed / expected IR between ER-associated genes vs other genes (background). The median fold change (arrow) is 1.68 (number of transcripts per group ER = 449, background = 8648).

Given the range of observed IRs we next asked whether the efficiency of initiation could be linked to gene function. Using gene set enrichment analysis we identified gene ontologies associated with extreme IR values (Fig. 4D, Table S2). Interestingly, genes with the highest IR were strongly enriched for membrane genes and other proteins synthesized exclusively at the endoplasmic reticulum (ER). Overall, these genes exhibited more CDS translation relative to their abundance of scanning compared to genes translated in the cytosol. To investigate whether this increased initiation is simply a consequence of better start codon contexts, we calculated their observed IR relative to the expected IR given the initiation sequence. This revealed a significantly higher IR compared to other genes with identical initiation contexts (Fig. 4E, median enriched 1.68 fold, p < 0.009) suggesting that these genes benefit from an overall increased initiation rate unrelated to the initiation sequence.

## Discussion

In this study we expanded the TCP-seq protocol in two key aspects: 1) we capture all small subunits, not only those that co-occur on transcripts with 80S ribosomes, and 2) we use template-switching in the library preparation to enable the use of less input material ^9^. Our modified RCP-seq therefore captures ribosomal complexes globally from all stages of the translation process and can be easily applied to other systems with limited input material, such as specific polysomal fractions or cell types. We used RCP-seq to study the dynamics of translation initiation during early stages of development in a vertebrate system, zebrafish. The longer 5’ UTRs of zebrafish allows for a detailed analysis of initiation by spatial separation of recruitment, scanning and start codon recognition.

Our data supports the threading model of ribosome recruitment to mRNA. At the 5’ end of mRNAs we observed a “ladder” of differentially sized fragments with 5’ ends coinciding with the transcription start site (Fig 2B). Fragment sizes shorter than the length of the 40S mRNA tunnel are consistent with the mRNA gradually entering the tunnel, but conflicts with a slotting model where single-sized fragments would be expected (Fig 2A). However, two alternative explanations could also potentially account for these fragments. In the first, the SSU could be slotted adjacent to the 5’ cap, but then proceed to back-slide in the 5’ direction. However, previous studies have shown that mRNA-binding by factors eIF4A, eIF4B/H and eIF4F prevent the SSU from back-sliding ^38,39^, which makes it unlikely that the abundant 5’ reads (suggesting a frequent occurrence) are due to back-sliding. The second possibility is that these reads are simply 3’-to-5’ degradation intermediates. However, two observations argue against this possibility: first, non-coding RNAs have very few 5’ reads arguing for a translation-dependent origin (Fig. S3G), and second, sequencing reads from degradation intermediates (and other possible artifacts) would be expected to increase when ribosome scanning is inhibited. Instead, upon eIF4E inhibition we observe that these short 5’ fragments disappear together with fragments derived from SSU scanning (Fig. 2C). We therefore conclude that threading of mRNAs is the most likely explanation for the presence of these fragments. Moreover, TCP-seq libraries from yeast realigned to CAGE-defined transcription start sites revealed a similar distribution of short 5’ fragments (Fig. S5) supporting threading as a universal mechanism,

Our data allowed to analyze the global processivity of scanning ribosomes. We found that the majority of scanning ribosomes reach the protein coding start codon and identified uORFs as a major cause of detachment. Consistent with previous studies ^25,40,41,42^, we find that the choice of stop codon affects uORF termination across the transcriptome, and furthermore that poor stop codons lead to an increase of read-through 80S ribosomes, decreasing the ability of SSUs to reinitiate at the CDS. This results in a reduction of CDS translation demonstrating that the choice of uORF stop codon can globally affect protein expression.

Finally, by accounting for the number of small ribosomal subunit complexes available for initiation, we confirmed previous observations that the Kozak sequence provides a strong initiation context. Nevertheless, our data revealed that there are additional endogenous sequences that give rise to equal or better rates of initiation. This is consistent with reports of weaker than expected correlations between Kozak sequences and gene expression in human ^36^ and yeast ^37^. We furthermore found that genes with the most efficient initiation rates were strongly enriched for proteins synthesized in the ER. While mRNAs at the ER have been previously shown to be more efficiently translated than mRNAs in the cytosol ^43,44^, our results further show that this increased translation is independent of the initiation sequence, but rather a result of SSU complexes proceeding more efficiently to translation elongation at the ER. Together, this shows that the IR metric can capture regulation of initiation in subsets of mRNAs and reveal novel features of compartmentalized translation.

Overall, our approach enables the deconvolution of translation initiation into distinct substeps. The RCP-seq protocol can be further applied to study samples with limited input material, which will allow addressing heterogeneity and specialization of the translation machinery, compartmentalized translation or tissue-specific translation. This opens for the possibility to obtain novel insights into scanning and initiating mechanisms across organisms and disease models.

## Methods

### Embryo sample collection and crosslinking

*D. rerio* embryonic samples were collected using standard zebrafish husbandry ^45^. In short, AB males and females were separated the day before mating. Shortly after first light, fish were put together and allowed to mate for 10 min. Embryos were collected in E3 medium (5 mM NaCl, 0.17 mM KCl, 0.33 mM CaCl_2_, 0.33 mM MgSO_4_), cleaned, and dechorionated using pronase (1 mg/ml, Sigma) for 5 minutes. Embryos were cleaned thoroughly after dechorionation and grown on 1% agarose plates containing E3 medium until the desired stage. Embryos were staged-matched for sample collection (Table S3). 200 embryos were transferred to 2 ml Eppendorf tubes per sample and washed twice in PBS with protease inhibitors (1:100 dilution, cOmplete, Mini, EDTA-free protease inhibitor cocktail) immediately prior to crosslinking, and left in 250 μl.

Embryos were snap-chilled by addition of 750 μl ice-cold PBS with 4% paraformaldehyde (PFA, freshly prepared) and immediately placed on ice. Samples were incubated for 15 min on ice, with gentle agitation. PFA medium was fully removed and 1 ml lysis buffer (20 mM HEPES-KOH pH 7.4, 100 mM KCl, 2 mM MgCl_2_) added. Glycine was added to a final 0.25 M concentration for PFA quenching, and samples incubated for 5 min on ice. Embryos were then washed twice in lysis buffer and resuspended in lysis buffer supplemented with 0.5 mM DTT, 2 μl Superase-In (RNase Inhibitor) and 1x protease inhibitor. Samples were immediately flash frozen in liquid nitrogen and stored at −80°C until used.

### Separation of ribosomal small subunit and ribosomal complexes

Samples were lysed in a cold room (4°C) by first shaking at 1,300 rpm for 10 min, followed by passing six times through a 27G needle. Samples were clarified by centrifuging for 15 min at 14,500g, 4°C. The OD260 absorbance of the supernatant was measured by nanodrop. 1/10th of each lysate was kept for RNA sequencing of that sample. Based on the absorbance readings, samples were digested using RNase I (0.0383 U × sample volume (μl) × OD 260 absorbance) for 45 min, at 23°C, 300 rpm shaking. Linear sucrose gradients were made from 5% to 30% sucrose solutions (containing 50 mM Tris-HCl pH 7.0, 50 mM NH_4_Cl, 4 mM MgCl_2_, 1 mM DTT) with Biocomp Gradient Station (long cap programme, 5% to 30%). Gradients were cooled down for 45 min at 4°C. Samples were layered on top of each gradient and tubes were centrifuged in a SW-41 rotor (Beckman-Coulter) at 4°C, 38,000 rpm for 4 hours. The gradients were fractionated using Biocomp Gradient Station and the small subunit and 80S fractions were identified and collected by monitoring the absorbance profile at 254 nm.

### RNA isolation

The collected fractions and RNA controls were supplemented with 1% SDS, 10 mM EDTA, 10 mM Tris-HCl pH 7.4, and 10 mM glycine. One volume of phenol:chloroform.isoamyl alcohol (pH 4.5) was added to each sample, immediately placed on a shaker at 65°C, 1,300 rpm for 45 min. After a 5 min centrifugation at 15,000g at room temperature, the aqueous phase was transferred to a new tube and precipitated by addition of 20 μg glycogen, 0.1 volume 3M sodium acetate (pH 4.5) and 2.5 volumes absolute ethanol. Samples were precipitated at −20°C for at least 3 hours. RNA was pelleted by centrifugation at 21,000g for 40 min, followed by two washes with 80% ethanol and 20 min centrifugation at 21,000g, 4°C. After drying the pellet, it was resuspended in 17 μl water, and concentration assessed on nanodrop.

### Construction of RCP-seq sequencing libraries

RNA samples were end-repaired by first incubating for 2 min at 80°C and on ice for 5 min, followed by the addition of 2 μl 10x T4 PNK buffer, 1 μl SUPERase-In, and 1 μl T4 PNK (10U/μl), and incubated for 2 h at 37°C. rRNA fragments were removed by using Ribo-Zero Magnetic Gold Kit (Illumina) following manufacturer’s instructions and purifying RNA using clean & concentrator-5 spin columns (Zymo Research) in a final volume of 13 μl. Libraries were constructed using the TaKaRa SMARTer smRNA-Seq Kit for Illumina, following manufacturers’ instructions without intermediate freezing points. In general, ATP was used for polyadenylation depending on the amount of starting material (less than 25 ng, no ATP). The number of cycles used was optimized per sample in the PCR amplification step and samples were eluted in a final volume of 20 μl. RCP-seq library sizes were checked on Agilent Bioanalyzer DNA High Sensitivity chips. Depending on size distribution, small and/or large fragments were removed by using AMPure XP beads (in order to remove adapter dimers and too large fragments not resulting from ribosomal protection). Sequencing was performed on a NextSeq 500 (Illumina) at the Norwegian Sequencing Centre in Oslo, high output mode with single reads of 75 bp or 150 bp.

### eIF4E inhibitor assays

The eIF4E inhibitor 4Ei-10 (or 6a ^12^) was synthesized at the Wagner lab (University of Minnesota, USA). The inhibitor was diluted in DMSO to a concentration of 100 μM, and kept frozen as stock. Further dilutions were performed in water. One nl of two concentrations (10 μM and 100 nM) were injected into dechorionated zebrafish embryos between 1-4 cell stages, in parallel with DMSO injections as controls. Embryos were allowed to continue development and samples were collected for RCP-seq at 64-cell and Shield stages (as described above). Flash frozen samples at Shield stages were also collected for polysome profiling in order to quantify effects on global translation for both 4Ei-10 and DMSO injected samples.

### Polysome profiling

Control (DMSO-injected) and inhibitor-injected samples were collected at Shield stage and flash frozen to halt ribosomes. Samples were lysed in polysome lysis buffer (10 mM Tris-HCl (pH 7.4), 5 mM MgCl_2_, 100 mM KCl, 1% Triton X-100) by shaking for 10 min at 1,300 rpm and 4°C, followed by 6x lysing through a 27-gauge needle. Samples were centrifuged at 14,500 g, for 15 min, 4°C. The supernatant was transferred to a new tube and its absorbance at 260 nm was measured with Nanodrop One to quantify RNA content. Linear sucrose gradients (15-45%) were prepared in a buffer containing 20mM HEPES KOH (pH 7.4), 5 mM MgCl_2_, 100 mM KCl, 2 mM DTT and 10 μl Superase-In. Gradients were cooled down for 45 min at 4°C. Samples were layered atop the gradients and ultracentrifuged in a Beckman-Coulter SW-41 rotor at 36,000 rpm, 2 h, 4°C. Polysome profiles of the gradients were obtained by in-line 254 nm absorbance measuring with Biocomp Gradient Station.

### Reporter assays

Translational efficiency of initiation contexts was tested using eGFP reporters. Three eGFP reporters with different initiation contexts were synthesized. The coding sequence of eGFP was amplified from pCS2+-eGFP vector using High Fidelity Phusion MasterMix (ThermoFisher, #F-531L). The forward primers for the PCR included the SP6 promoter sequence, followed by 26 bp of eIF3d leader sequence and the respective start codon context; the reverse primer (M13-rev) was located after the common SV40 termination signal included in the pCS2+ vector backbone (see Table 1). Following synthesis by a two-step PCR, samples were gel-purified and RNA was synthesized with SP6 mMessage mMachine (Thermo Fisher, #AM1340), following manufacturer’s recommendations, and cleaned-up using Zymo RNA Clean & Concentrator-25 columns (Zymo Research, #R1013).

**Table 1:**
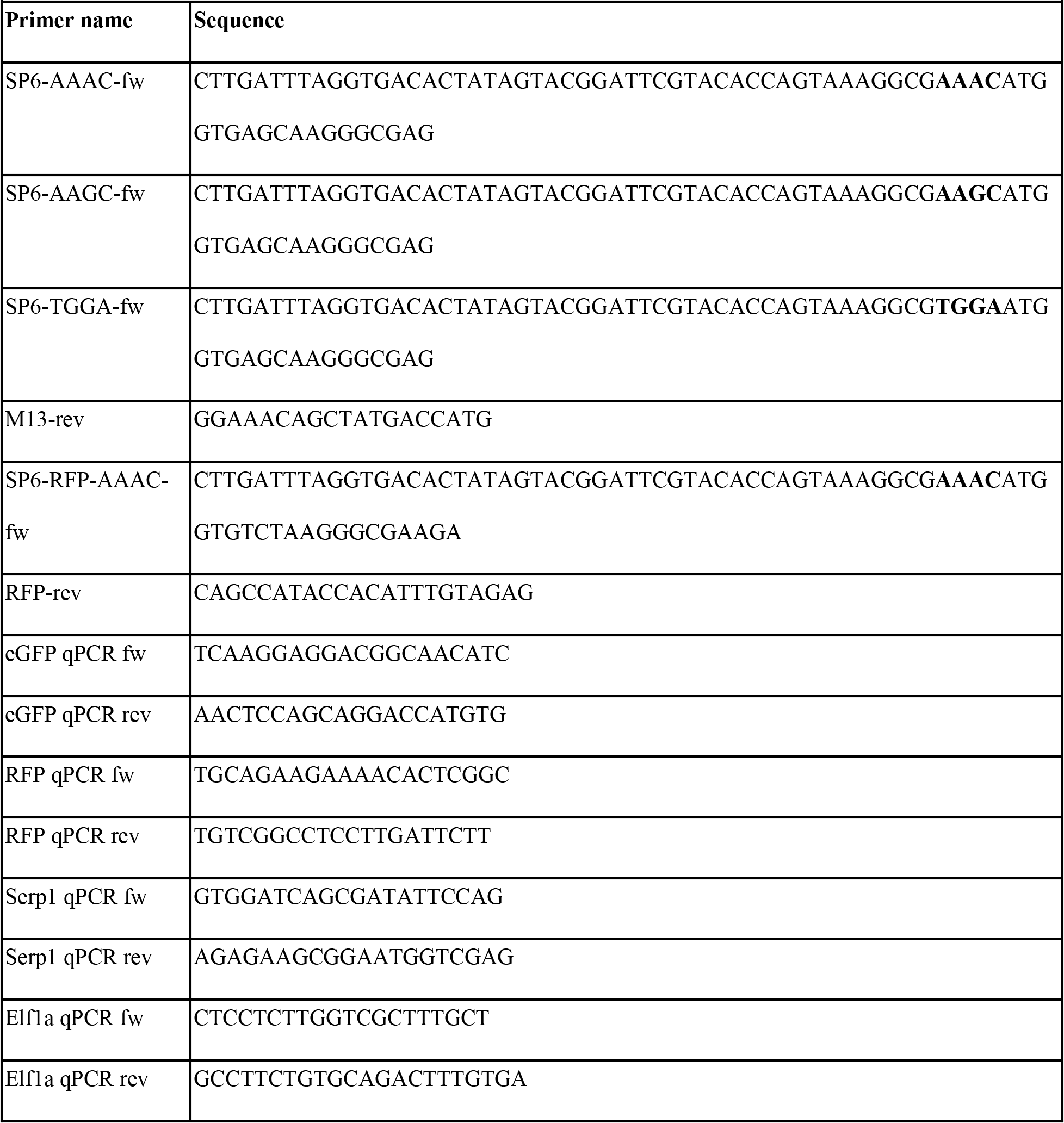
primers used for reporter synthesis and qPCR

To control for GFP expression changes, we further synthesized RFP mRNA with fixed start codon context. The coding sequence of RFP was amplified from pT2KXIGdeltaIn-MCS-birA-tagRFP vector (AddGene #58378) using High Fidelity Phusion MasterMix. The forward primers for the PCR included the SP6 promoter sequence and the reverse Primer was located after the common SV40 termination signal included in the pT2KXIG vector backbone (see Table 1). Following synthesis by a two-step PCR, samples were processed as above to obtain mRNA.

Stock RNA solutions for injections at 120 ng/ul were made based on nanodrop & Qubit RNA concentration measurements. RNA solutions containing RFP and eGFP reporters were co-injected at a final concentration of 50ng/ul per reporter (with phenol red added for injection visualization) and injected at 1 nl per embryo at 2-4 cell stage (dechorionated embryos). Embryos were collected and dechorionated as described above. Groups of 25 embryos for each of the samples were collected at 24 hpf and used for qRT-PCR and eGFP protein quantification. For qRT-PCR, RNA was extracted using Trizol, and processed following the protocol previously described ^46^. eGFP RNA expression was quantified against RFP for injection control and endogenous controls *serp1* and *Elf1a*.

For eGFP protein quantification, the 25 embryos collected were homogenized in 100 mM Tris (pH 7.5) with 1% Triton X-100, and processed as described in ^34^. After quantifying total protein concentration with the Pierce BCA kit (Thermo Scientific), the samples were diluted to equal total protein concentrations. A Qubit fluorometer (Life Technologies) was used to measure eGFP fluorescence, using 9 technical replicates for each sample, and with GFP dilutions as measurement controls.

Translational efficiency was calculated as described previously ^34^, by quantifying RNA with qRT-PCR and quantifying eGFP protein by measuring eGFP fluorescence in Qubit. Relative eGFP protein (average of all 9 technical replicates for each sample) was calculated compared to AAAC (zebrafish Kozak), divided by the relative abundance of eGFP RNA in each corresponding sample as measured by normalised qPCR abundance. For the AAAC samples, we compared their eGFP values to the average of the group to get their individual TE values.

### Transcript definitions

The analysis was performed on the most highly expressed transcript from each gene, calculated from total RNA-seq coverage (datasets are described in Table S4). Cap analysis gene expression (CAGE) was used to update the 5’ UTR on a per sample basis as follows. The highest CAGE peak was selected in a search region from the 3’ most end of the transcript, to the greater of; the 5’ most end of the 5’ UTR or 1000 nt upstream of the annotated start codon (If the highest CAGE peak was called downstream of the protein TIS the transcript was excluded for further analysis). Transcripts were excluded from further analysis if they overlapped with the most highly expressed transcript of another gene, or with an annotated non-coding transcript (defined as all Ensembl transcripts with biotypes other than protein_coding).

TISU motif containing transcripts were defined as those that with 5’ UTRs of <= 30 nt in length and a PWM scores against the consensus TISU sequence SAASATGGCGGC (where S is C or G) of >= −12. TOP motif containing transcripts were defined as those beginning with a C followed by at least 4 T nucleotides. Kozak sequence strength was determined through PWM scores against the zebrafish kozak matrix taken from ^34^. Three genes (ENSDARG00000077330, ENSDARG00000102873, ENSDARG00000089382) contained strong coverage peaks across repetitive regions in their 3’ UTR and were excluded from plots showing densities over 3’ UTRs (Fig. 1B, S14).

### Read trimming and alignment

RCP-seq reads were trimmed with cutadapt searching for the “AAAAAAAAAA” added to the 3’ of each fragment during library preparation, allowing 1 mismatch and at least 5 nt of overlap. The first 3 nt of each read were removed, reads shorter than 15 nt after trimming were discarded. The remaining reads were aligned to, in order, the PhiX genome, rRNA from the silva database^47^ (version 119), organism-specific ncRNA as defined by Ensembl (zebrafish GRCz10), organism-specific tRNA produced with tRNA-scan SE ^48^ (default settings). Reads that did not match to any of the above were aligned to the *D. rerio* GRCz10 genome. Total RNA-seq reads were trimmed and aligned to the *D. rerio* GRCz10 genome. Ribo-seq reads were trimmed and aligned to rRNA and ncRNA as above, unaligned reads where then aligned to the *D. rerio* GRCz10 genome. CAGE reads were trimmed and aligned to the *D. rerio* GRCz10 genome. Alignments were performed using tophat2 ^49^ against the *D. rerio* GRCz10 genome and ensembl version 81 gene annotations (Table S5), reporting up to 20 hits for reads mapping to multiple locations (later filtered with MAPQ, see below).

TCP-seq data from *Saccharomyces cerevisiae* ^8^ (SRA: SRP074093) was processed as above (with the exception of 1st 3nt removal and using STAR as aligner with default parameters) to the R64_1_1 genome with Ensembl version 79 gene annotations. *S. cerevisiae* 5’ UTR were defined with CAGE ^11^ (GEO: GSE69384).

### RCP-seq fractionation

The RCP-seq sedimentation fractions corresponding to small subunits and 80S complexes were determined from sequencing all sedimentation fractions. Based on coverage profiles (Fig. S14), fractions 12-14 were determined to contain small subunit fragments. Fractions 18-19 were determined to contain 80S complex fragments. RCP-seq small subunit counts in this study are reported from a pooled set of all relevant fractions (Table S3), unless otherwise stated. In order to determine the maximum length of small subunit RCP-seq reads, one sample was re-sequenced for 150 cycles (shown in Fig. S4), as opposed to the 75 cycles used for the rest of the samples.

### Further read processing

Inappropriately truncated RCP-seq reads after polyA trimming were updated by extending alignments where trimmed regions exactly matched transcriptomic references. Unusually high RCP-seq coverage peaks were removed from transcripts by filtering out reads with the same 5’ and 3’ coordinates that were present at >=200 times the average coverage of each transcript. A subset of small subunit reads (~25-35 nt in length), were observed to show 3nt periodicy over the CDS region (Fig. 1D, lower left, CDS region). This periodicity is indicative of translation, but it was not clear if these reads represent the leaky scanning of 43S PICs, queued behind translating ribosomes, or footprints of translating complexes, where possibly the 60S subunit has become detached, before sedimentation. As such, reads corresponding to the length of typical translating fragments (length 25-35 nt) were considered ambiguous and removed from RCP-seq small subunit libraries. Conversely the RCP-seq 80S libraries used in figure 1B, were filtered to only include the lengths of typical translating fragments (length 25-35 nt), akin to ribo-seq libraries. RCP-seq reads that mapped to positions overlapping the last 10 nt of each transcript were discarded, to remove 3’ peaks that likely result from polyA selection during the library preparation.

### Read counting

The relative enrichment of tRNA species between small subunit and 80S complex fractions was calculated with edgeR ^50^ using a binomial generalized log-linear model and likelihood ratio test. Read counts were summed per tRNA anticodon type. CAGE reads counts were normalised using the power law method from the cageR package ^51^. The FPKMs (fragments per kilobase, per million mapped reads) of RCP-seq, Ribo-seq and Total RNA-seq reads with a MAPQ >= 10 were calculated for transcript 5’ UTR, CDS, and 3’ UTR regions. Reads that overlapped multiple regions were preferentially assigned to CDS > 5’ UTR > 3’ UTR.

### Fragment lengths heatmaps and fragment distribution

Metaplots of RCP-seq footprint distributions were plotted in windows across at the 5’ most end of transcripts (TSS) or around protein TIS, for transcripts with at total RNA-seq FPKM>10 and 5’ UTRs at least 100 nt long. Fragment counts are assigned to either the 5’ or the 3’ end of the fragment. The heatmaps of counts per fragment length are coloured by sum of counts from all transcripts at a given position for a given fragment length. The proportion of coverage window counts are summed for all transcripts at given position. The transcripts of genes ENSDARG00000036180, ENSDARG0000001479 were observed to have strong artefactual peaks caused by premature read trimming in polyA regions upstream of the protein TIS and were removed from the TIS plots in Fig. 1D.

For the yeast TCP-seq data, the footprints were similarly plotted as heatmaps for the area around the TSS in Fig. S5. Using a median based filter to remove transcripts that had extreme peaks in the +2 to +40 region relative to TSS (total of 21 genes). Filter removed transcript if: peak at any position in +2 to +40 > median footprints in transcript per position * 99 quantile rank of transcript’s footprints per position + 99.9 quantile rank of the medians of all transcripts.

### Scaled coverage meta plots

The coverage, or 5’ counts of RCP-seq and ribo-seq reads with a MAPQ >= 10 was calculated across transcript 5’ UTRs, CDS, and 3’ UTR. Transcripts with 5’ UTRs, CDS, and 3’ UTR greater than a length cutoff (typically 100 nt) were scaled to the length value. Values are displayed as the sum of all selected transcripts, mean normalised values or z-score for all selected transcripts. Counts from each transcript normalised by z-score across the whole transcript allow for comparisons of transcripts across wide expression ranges.

### Estimates of 43S PIC loss across 5’ UTRs

The coverage of small subunit footprints in regions proximal to the beginning of the transcript and the protein TIS were used to infer the number of 43S PICs recruited to a transcript and those available for initiation at the protein TIS. These regions were defined based on coverage metaplots (Fig. S10) as +70 to +120 nt relative to the beginning of the transcript and −100 to −50 nt relative to the protein TIS. The ratio of coverage in these regions was used to estimate the loss of 43S PICs across 5’ UTRs for all protein coding transcripts, with RNA-seq FPKM>= 10 and 5’ UTR >= 220 nt in length.

### 5’ feature plots (Fig. 2)

Empirical cumulative density for scanning efficiency (Fig. 2D,E) and translational efficiency (Fig. S9) were plotted for all protein coding transcripts with >= 10 RNA FPKM. For groups of transcripts starting with; **i**) an A, C, G or T; or **ii**) transcripts starting with a TOP motif, transcripts starting without a TOP motif and also starting with a C, and transcripts starting without a TOP motif and also starting with a A, G or T.

### uORF plots (Fig. 3)

Upstream open reading frame coverage metaplots were produced for all protein coding transcripts with an ATG uORF starting > 100 nt from the 5’ most end of the transcript and the TIS, centering on the first (5’ most) ATG uORF within the transcript.

The proportion of small subunit reads mapping upstream (from the beginning of the transcript to the uORF start codon) or downstream of the uORF (uORF start codon to the protein TIS) were calculated for all ATG, CTG and GTG uORFs, starting >= 50 nt from the beginning of the transcript and the protein TIS, in transcripts with RNA FPKM>=1, stratified by uORF start codon, and Kozak strength quantile. Similarly the proportion of small subunit reads mapping upstream (from the beginning of the transcript to the uORF start codon) or downstream of the (uORF stop codon to the protein TIS) were calculated for all ATG, CTG and GTG uORFs starting >= 50 nt from the beginning of the transcript and >=50 nt from uORF stop codon to the protein TIS, in transcripts with >=1 RNA FPKM, stratified by uORF start codon, and uORF stop codon.

Scanning efficiency, translational efficiency and 5’ UTR translational efficiency (Fig. S8, S15) were calculated for all protein coding transcripts with 5’ UTRs >= 100 nt in length and RNA FPKM>=1, that contained 0,1,2,3 or 4 ATG uORFs. Statistical significance was reported between transcripts containing 0 ATG uORFs versus those containing 1-4 ATG uORFs as: W = 112420000, p-value < 2.2×10^−16^ for 5’ UTR RCP-seq 40S FPKM, W = 138210000 p-value < 2.2×10^−16^ for translational efficiency and W = 73953000, p-value < 2.2×10^−16^ for 5’ UTR translational efficiency.

The ratio of RCP-seq small subunits to ribo-seq 80S complexes over the uORF stop codon, per uORF stop codon; the density of 80S complexes between the uORF stop codon and protein TIS, normalised by CDS total RNA-seq; and the translational efficiency (Fig. S12) were calculated for all ATG, CTG or GTG uORFs starting >= 50 nt from the beginning of the transcript and >=50 nt from uORF stop codon to the protein TIS, in transcripts with >=1 RNA FPKM. Statistical significance was reported as the median log2 ratio of small subunits to 80S over the stop codon of transcripts containing TAA uORFs vs TGA uORFs (W = 250820000, p-value < 2.2×10^−16^ the median normalised log2 downstream ribo-seq 80S density of TAA uORFs vs TGA uORFs (W = 1190600000, p-value < 2.2×10^−16^), and the median log2 translational efficiency of transcripts containing TAA uORFs vs TGA uORFs (W = 1298200000, p-value = 6.128×10^−11^).

### Initiation plots (Fig. 4)

Initiation rates (see Fig. S8) were calculated as the ratio of RCP-seq small subunits in the 5’ UTR to ribo-seq 80S complexes in the CDS, for all protein coding transcript sequences with RNA FPKM >= 10 and 5’ UTRs >= 100 nt in length. Initiation rates for transcripts were then grouped by each nucleotide, per position in a −4 to +5 window surrounding the protein coding start codon, excluding the start codon. Median initiation rate was calculated for nucleotides that were present more than 1000 times. Initiation rates were also calculated over continuous sequence contexts, using a smaller window of −4 to +3 nucleotides surrounding protein coding start sites, calculating the median initiation rate for all sequences present >= 20 times in the selected transcripts. This smaller window was used in order to increase the number of transcripts present in each sequence bin.

### GO enrichment analysis

GO analysis was performed with GOrilla, detecting GO terms based on ranking of genes according to Initiation Rate (IR). P-values computed according to the mHG model, and FDR q-values for the correction of the p-value for multiple testing. Genes were filtered by leader lengths >= 100nt, CAGE peaks on transcript > 5 reads, and FPKM of RNA-seq > 0 and FPKM of ribo-seq and SSU libraries > 0. Giving a total of 449 ER enriched genes and 8648 as the background set.

### Statistical testing and plotting

Significance testing was performed in R using the Wilcoxon rank sum test with continuity correction. The metrics used to investigate relationships between small subunits and 80S footprints are summarised in Fig. S15. Boxplot upper whiskers extend from the 1st quartile to the largest value no further than 1.5 times the distance between the first and third quartile, from the first quartile. Boxplot lower whiskers extend from the third quartile to the smallest value no further than 1.5 times the distance between the first and third quartile, from the third quartile.

## Data and code availability

Custom scripts used to process the RCP-seq libraries are available at the following link: https://github.com/agiess/RCP_processing. The RCP-seq libraries have been uploaded to the ENA database privately until publication (accession number PRJEB33323). Access will be granted upon request.

## Acknowledgements

We wish to thank members of the Valen and Thompson labs (University of Bergen, Norway), Ignatova and Duncan labs (University of Hamburg, Germany), and Preiss lab (Australian National University, Australia) for helpful discussions of methodologies. We wish to thank Andrea Pauli (IMP, Austria), Sushma Nagaraja-Grellscheid and Eric Thompson (University of Bergen, Norway) for providing valuable feedback on the manuscript.

The project was funded by Bergen Research Foundation, the Norwegian Research Council (#250049) and core funding from the Sars International Centre for Marine Molecular Biology.

## Author contributions

AG, YTC and EV designed the research. YTC, MK, TBB and SS performed the experiments. AG, YTC, HT and EV analyzed the data. AO and CRW designed and synthesized the 4Ei-10 compound. All authors discussed the results and contributed to writing the paper.

