## Supplementary files for "Deconstructing the individual steps of vertebrate translation initiation"

**This file includes:**

Supplementary Figures S1-S15

Supplementary Tables S1-S5

Supplementary Notes

### Supplementary Figures

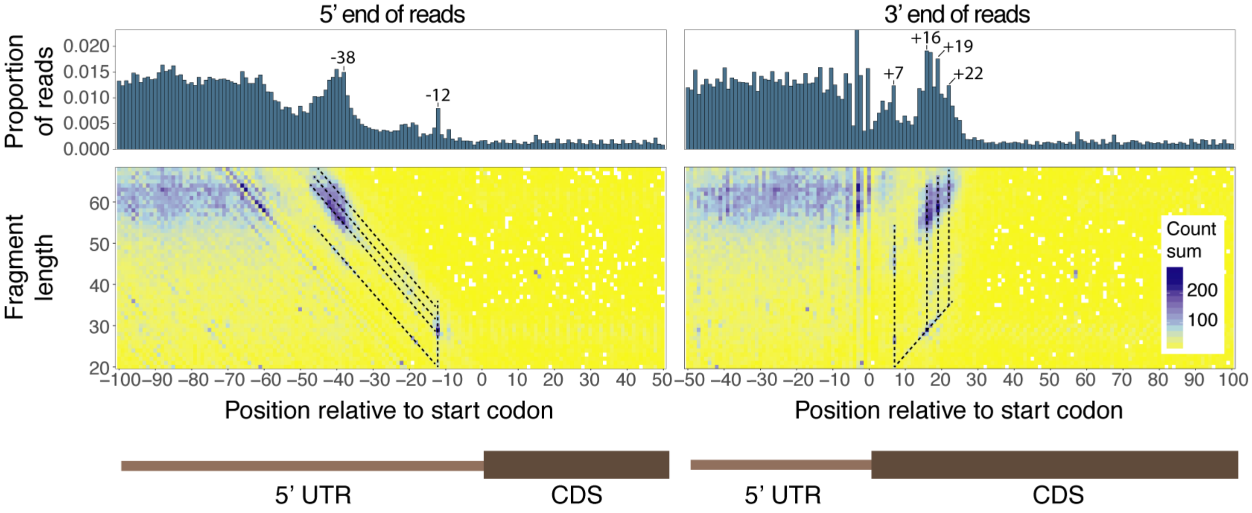


***Figure S1:*** **Danio rerio** ***read populations at transcription initiation sites.***

*Conformations of RCP-seq small subunit fraction footprints around start codons. Counts of 5' (left) or 3' (right) ends of fragments are summed across highly expressed genes (>= 10 FPKM). The barplots show the proportion of read counts per position (x-axis), while the heatmaps show the same counts stratified by length (y-axis) and coloured by total count. Black dotted lines highlight the major read populations, which are broadly consistent with the initiating read confirmations described in yeast, beginning at -12 and ending at +6, +16, and +24 nt (in relation to the start codon)* [1].

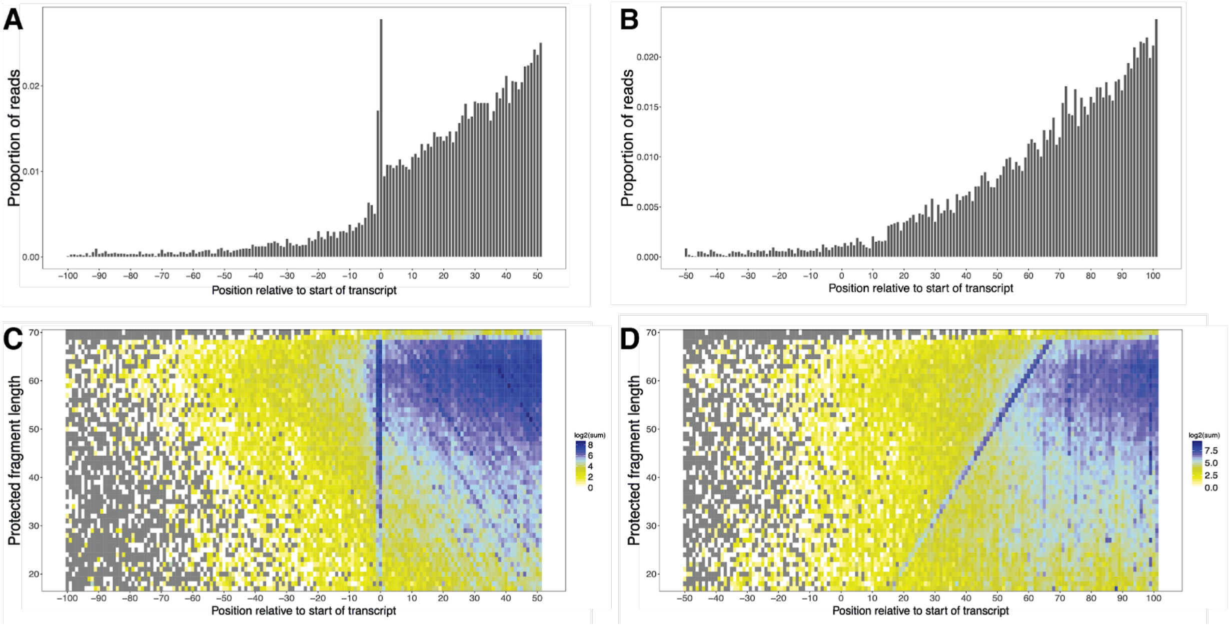


***Figure S2:*** **D. rerio *read density and fragment length metaplots at transcription start sites.***

*Metaplots of small subunit footprint distributions at 5' most end of* D. rerio *transcripts using counts assigned to the 5' (**A, C**) or 3' ends (**B, D**) of fragments. **A-B**) The barplots show the proportion of read counts per position (x-axis) relative to transcription start site, while the heatmaps (**C-D**) show the same counts stratified by length (y-axis) and coloured by total count (represented as log2).*

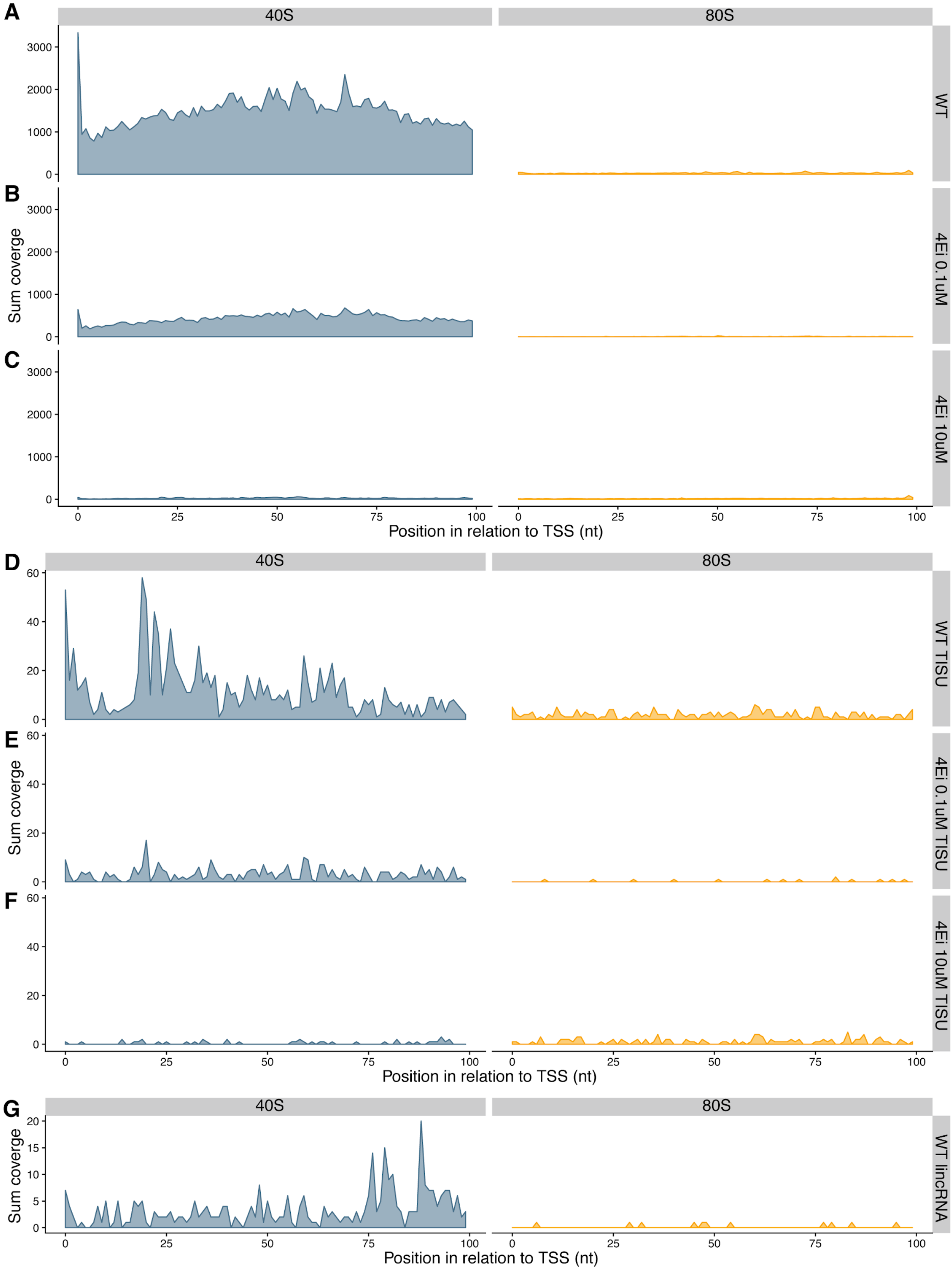


***Figure S3: 5' read count metaplots at transcription start sites.***

*Transcript 5' read count metaplots of 64-cell* D. rerio *RCP-seq small subunit footprints (blue) and 80S ribo-seq (orange), in transcripts with 5' UTR regions >= 100 nt. Showing the first 100 nt of 5'UTRs, contributions from each transcript are summed.* ***A****) Protein coding transcripts of WT samples.* ***B****) Protein coding transcripts with 0.1 µM 4Ei-10 treatment.* ***C****) Protein coding transcripts with 10 µM 4Ei-10 treatment.* ***D****) Protein coding TISU transcripts of WT samples.* ***E****) Protein coding TISU transcripts with 0.1 µM 4Ei-10 treatment.* ***F****) Protein coding TISU transcripts with 10 µM 4Ei-10 treatment.* ***G****) lincRNA transcripts of WT samples. Protein coding transcripts (n=5861), lincRNA transcripts (n= 715), TISU transcripts (n=207).*

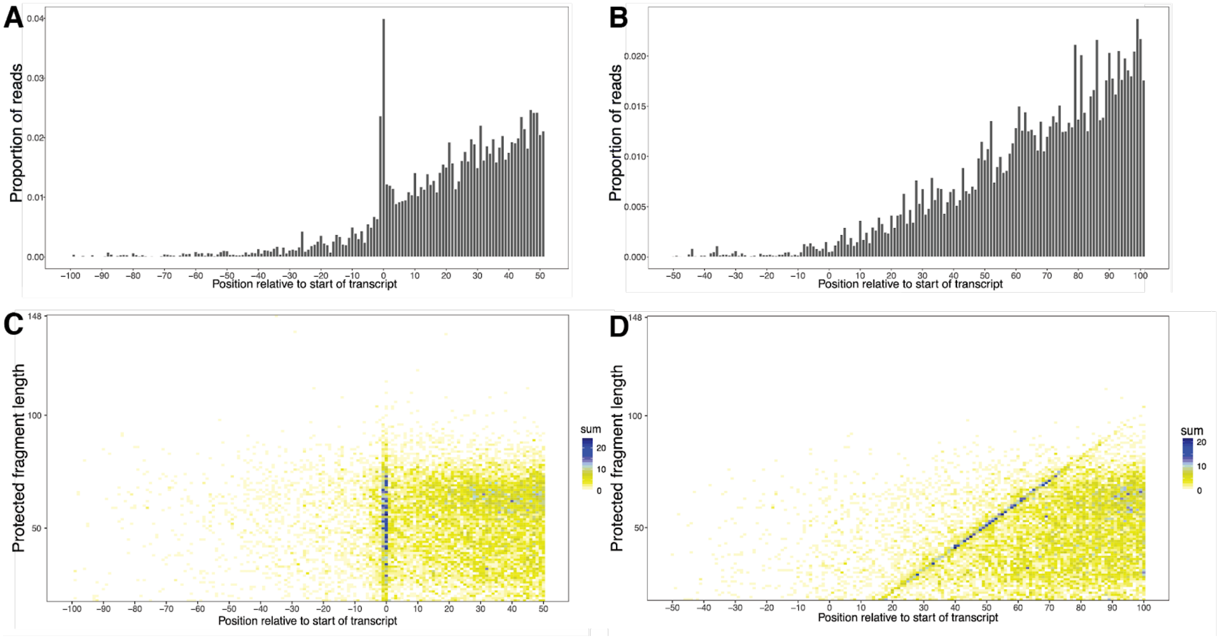


***Figure S4: Beginning of transcript density and fragment length metaplots of 150 nt sequenced library.***

*Metaplots of small subunit footprint distributions at 5' most end of* D. rerio *transcripts using counts assigned to the 5' (**A, C**) or 3' ends (**B, D**) of fragments. **A-B**) The barplots show the proportion of read counts per position (x-axis) relative to transcription start site, while the heatmaps (**C-D**) show the same counts stratified by length (y-axis) and coloured by total count.*

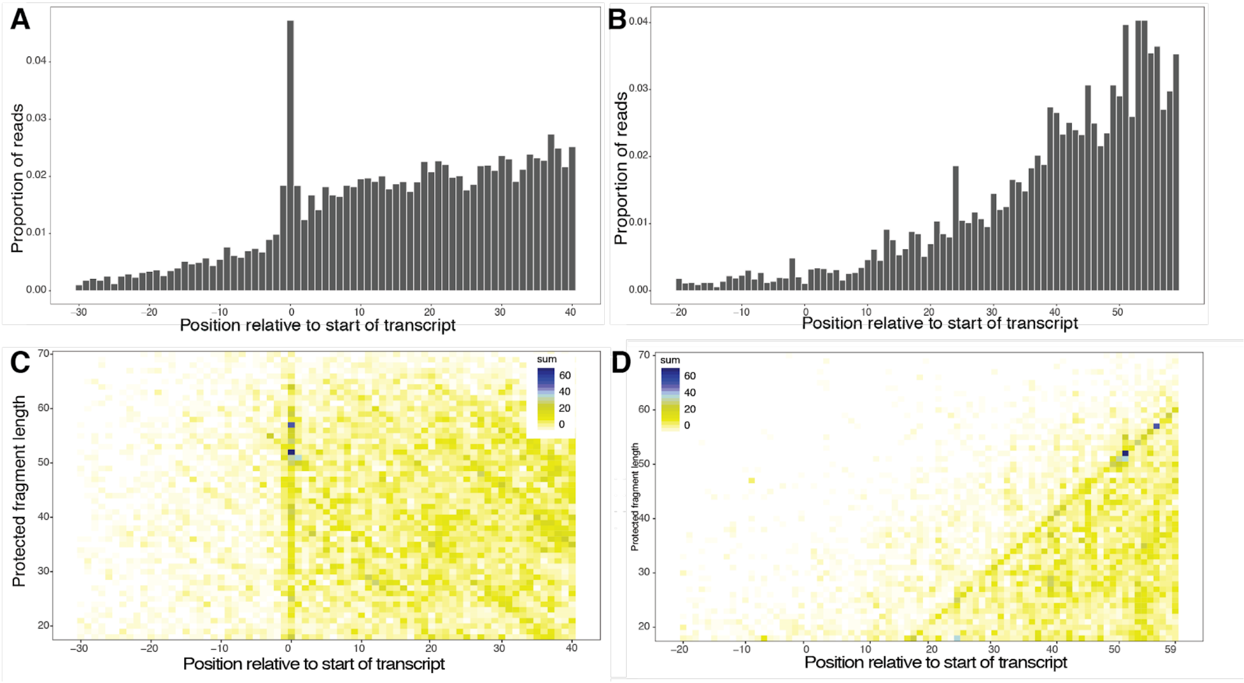


***Figure S5:*** **S. cerevisiae *read density and fragment length metaplots at transcription start sites.***

*Metaplots of small subunit footprint distributions from TCP-seq* [1] *at 5' most end of* S. cerevisiae *transcripts using counts of the 5' (**A, C**) or 3' ends (**B, D**) of fragments, using CAGE-defined transcript start sites* [2]. ***A-B****) The barplots show the proportion of read counts per position (x-axis) relative to transcription start site.* ***C-F****) The heatmaps show the same counts stratified by length (y-axis) and are coloured by total count (**E-F**). Genes with abnormal peaks were removed, a total of 21.*

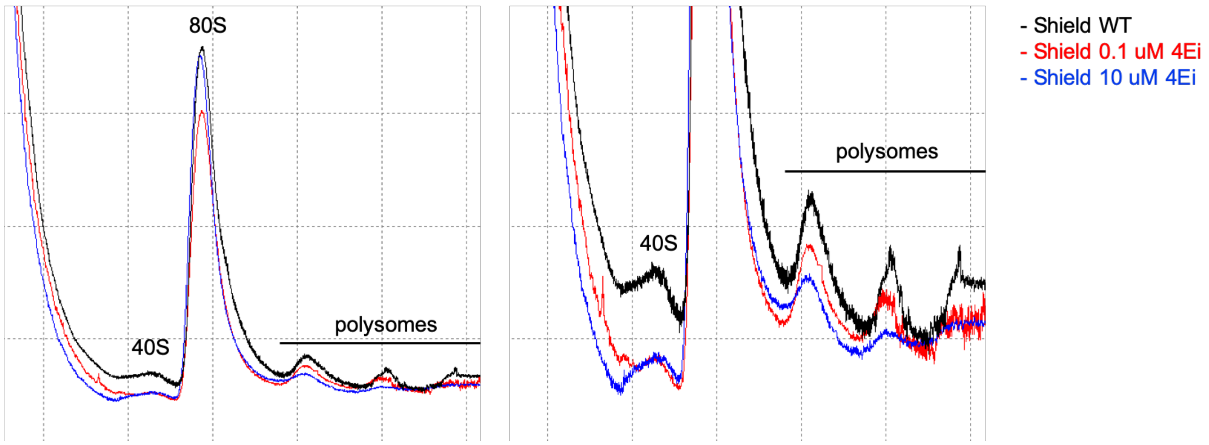


***Figure S6: General inhibition of translation following 4Ei-10 treatment.***

*Polysome profiles of zebrafish embryos at the Shield stage for WT and in the presence of eIF4E inhibitor at two different concentrations (0.1 µM and 10 µM). A dose-dependent reduction in polysomes is observed following 4E inhibition.*

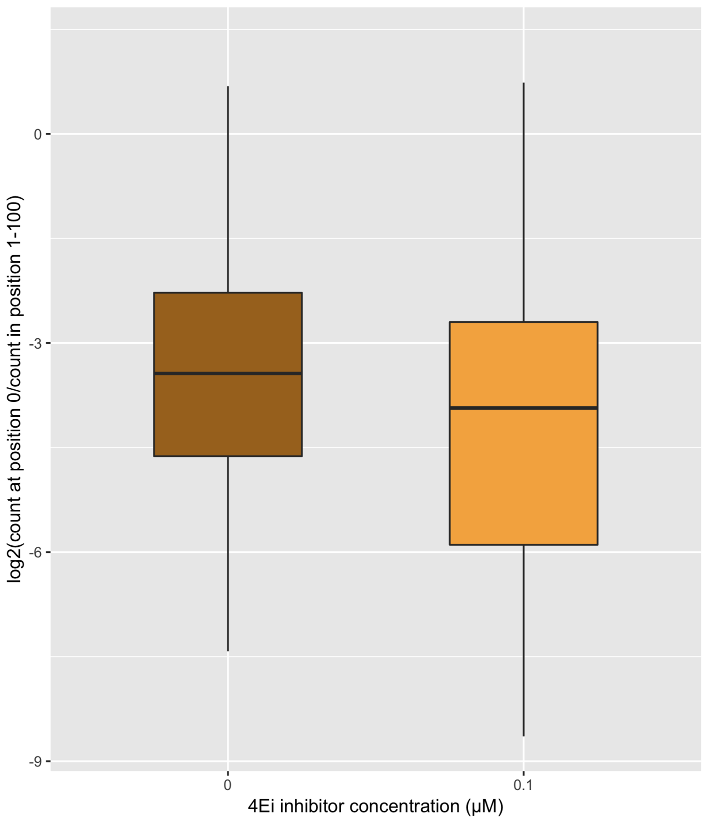


***Figure S7: Ratio of 5' read counts over 5' most position by reads in the following 99 nucleotides.***

*The ratio of 5' read counts at the transcription start site (first nucleotide) of each transcript, normalized by the 5' read count in position 2-100 in each transcript, in WT (0 μM) (brown), and 4Ei-10 (0.1 μM) treated (orange) samples. Transcripts were selected to have at least 5 reads in the first nucleotide position (n = 126) and a 5' UTR > 100nt.*

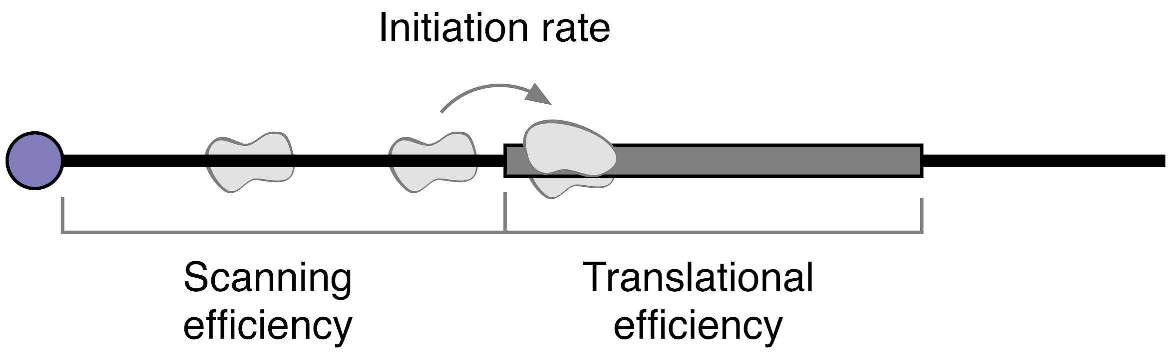


***Figure S8: Schematic representation of the three introduced metrics.***

*Scanning efficiency (SE) is defined as the number of small subunit footprints over a 5' UTR relative to its mRNA abundance. A higher SE indicates more 43S PICs per RNA. Initiation rate (IR) is defined as the ratio of 80S ribosomes in the CDS to small subunits in the 5' UTR. A higher IR indicates more translation per unit of scanning. Translational efficiency (TE) measures elongating ribosomes relative to mRNA abundance. A higher TE indicates more 80S ribosomes per RNA molecule.*

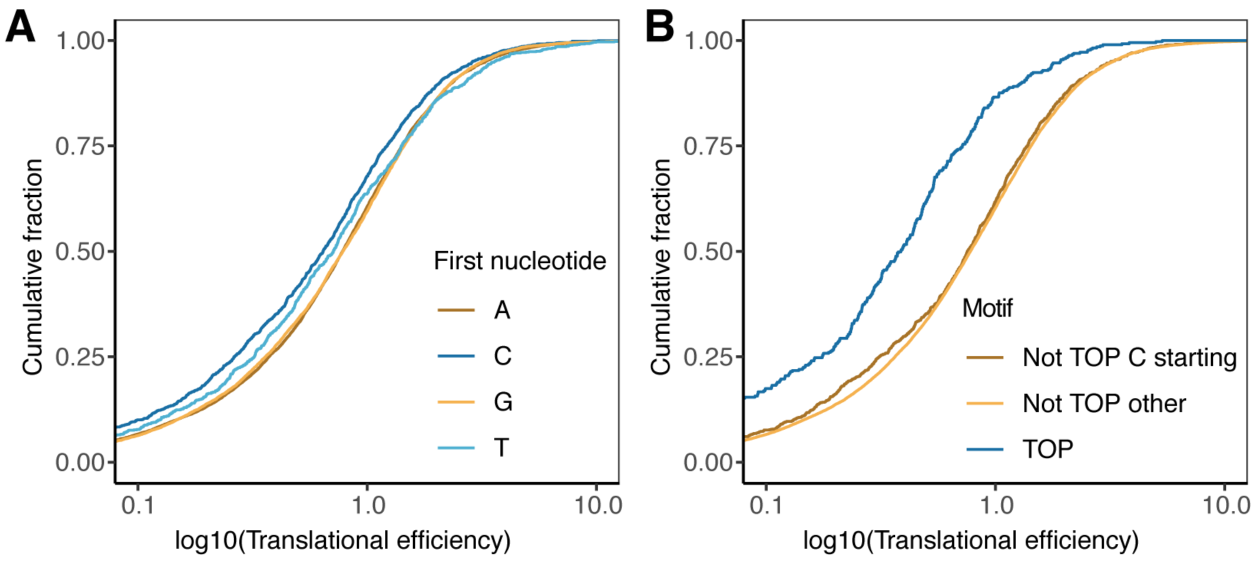


***Figure S9: 5' transcript features vs translational efficiency.***

*Empirical cumulative density of translational efficiency (TE) for highly expressed transcripts (>= 10 RNA FPKM). (**A**) Transcripts colored by their first nucleotide. An initial pyrimidine (C/T) results in lower TE than a purine (A/G) (C: $p < 5.8x10^{-16}$, T: $p < 0.0075$) (number of transcripts per group: A = 10388, C = 1663, G = 8335, T = 967). (**B**) Transcripts starting with a TOP motif (13 pyrimidines) show reduced TE ($p < 2.2x10^{-16}$) and non-TOP transcripts starting with a C to a smaller extent also show reduced TE (number of transcripts per group: Not TOP C starting = 1257, Not TOP other = 19690, TOP = 406).*

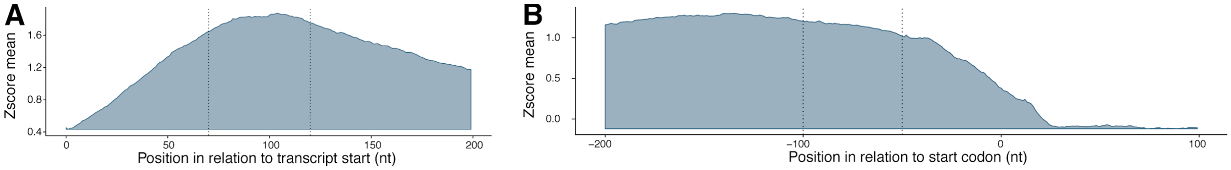


***Figure S10: Regions for estimations of small ribosomal subunit loss across 5' UTRs***

*Transcript read coverage metaplots of RCP-seq small subunit complex footprints (blue) expressed as the average over z-scores calculated for each transcript (emphasizing the shape of a typical mRNA distribution). The regions used for estimating small subunit densities at the beginning of the transcript (**A**) and proximal to the start codon (**B**) are indicated with dashed lines. Transcripts were selected to be protein coding with 5' UTR regions >= 200 nt.*

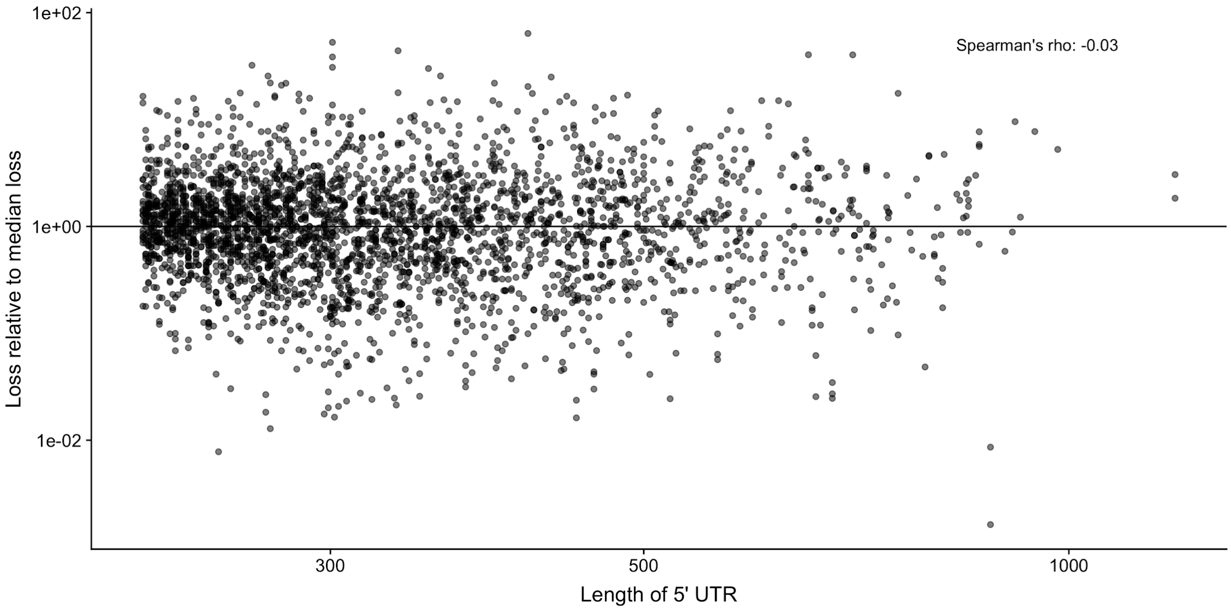


***Figure S11: Loss of small subunits scanning the 5'UTR as a function of 5' UTR length.***

*The loss of subunits across the 5' UTR (y-axis) as a function of 5'UTR length (x-axis). To control for the strong dependency between length of 5'UTR and number of uORFs, the loss is calculated relative to the median loss of small subunits for all 5'UTRs with the same number of ATG-initiated uORFs (horizontal line). Transcripts are selected to have 5'UTR > 220 nt and 5' proximal SSU > 10 FPKM and TIS proximal regions SSU > 0 FPKM.*

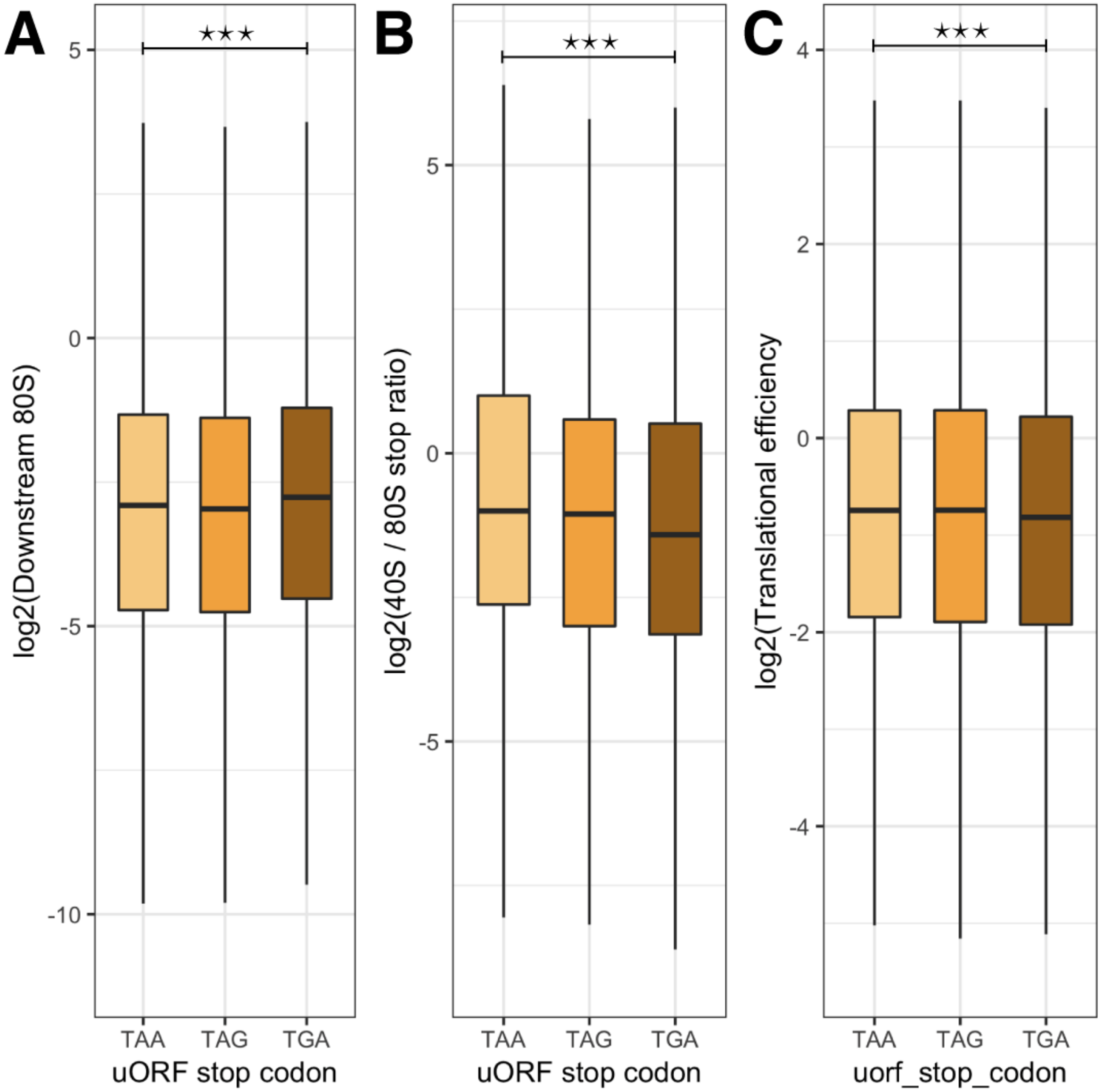


***Figure S12: uORF stop codons affect downstream translation.***

*The choice of stop codon in the uORF leads to different translational termination efficiency of downstream CDS, with TAA and TGA being the least and most repressive, respectively. Measured as (**A**) density of ribo-seq 80S complexes downstream of the uORF (and before the CDS TIS) (TAA uORFs −3.95, vs TGA uORFs −3.74, $p < 2.2x10^{-16}$), (**B**) the ratio of RCP-seq small subunit footprints to ribo-seq 80S complex footprints over uORF stop codons (TAA uORFs -2.18, vs TGA uORFs -4.25, $p < 2.2x10^{-16}$), and (**C**) effects on TE of downstream CDS (TAA uORFs -0.84, vs TGA uORFs -0.89, $p < 6.128x10^{-11}$). Number of transcripts per group TAA = 4619, TAG = 3170, TGA = 4603.*

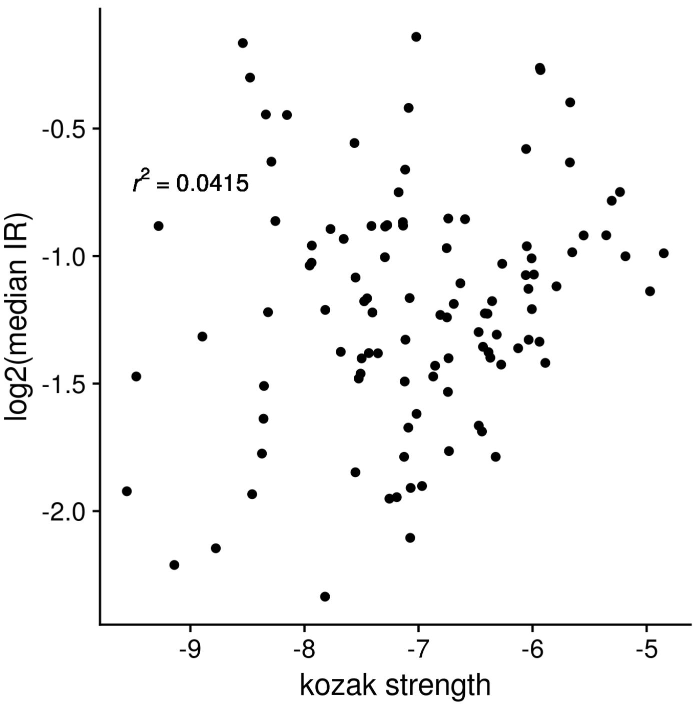


***Figure S13: Correlation between Kozak strength and Initiation Rate.***

*Numerical values for colors shown in Figure 4B. There is a weak positive correlation between Initiation Rate (IR) and similarity to the Kozak sequence. The sites with very high similarity to the Kozak (> -5.5) tend to have good IR, but overall Kozak similarity is not a good predictor of IR.*

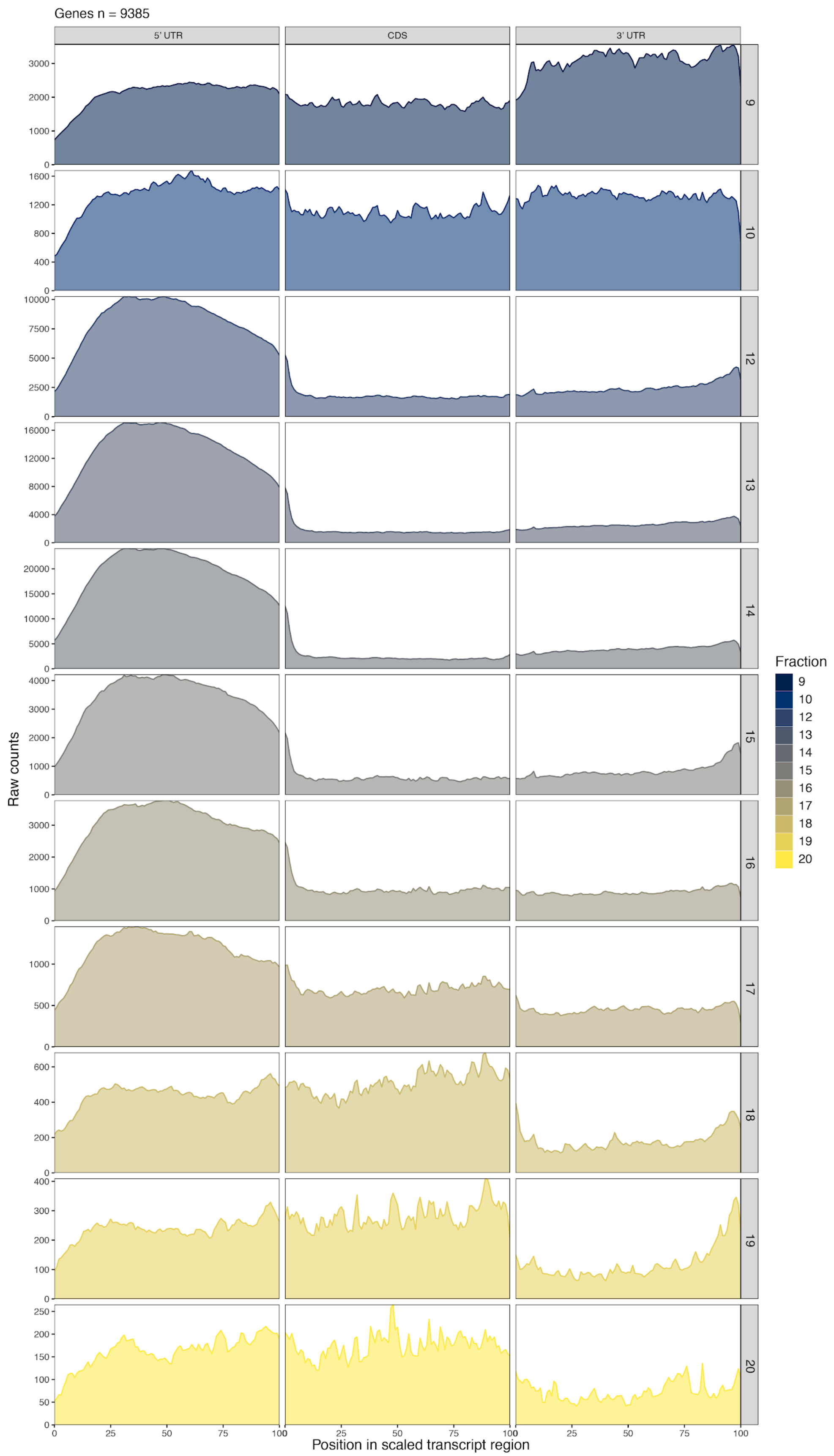

Genes n = 9385
5' UTR
CDS
3' UTR
Raw counts
Position in scaled transcript region
Fraction
9
10
12
13
14
15
16
17
18
19
20

***Figure S14: Coverage metaplots of all RCP-seq fractions.***

*Transcript coverage metaplots of separate fractions from RCP-seq sedimentation steps (see methods), contributions from each transcript are shown as the raw counts Fractions 12-14 were selected to represent the small subunit complex footprints and fractions 18 and 19 for translating 80S complex footprints.*

uORF small subunit (SSU) consumption rate for start codons:

$$\frac{SSU\ coverage\ upstream\ of\ uORF\ start\ /\ length\ of\ upstream\ region}{SSU\ coverage\ downstream\ of\ uORF\ start\ \ /\ length\ of\ downstream\ region}$$

uORF small subunit consumption rate for stop codons:

$$\frac{SSU\ coverage\ upstream\ of\ uORF\ start\ /\ length\ of\ upstream\ region}{SSU\ coverage\ downstream\ of\ uORF\ stop\ /\ length\ of\ downstream\ region}$$

uORF stop codon recognition rate:

$$\frac{SSU\ count\ at\ uORF\ stop\ codon}{80S\ count\ at\ uORF\ stop\ codon}$$

uORF readthrough rate:

$$\frac{80S\ count\ downstream\ of\ uORF\ stop\ codon\ /\ length\ of\ downstream\ region}{CDS\ RNA\ FPKM}$$

5’ UTR translational efficiency:

$$\frac{5'\,UTR\ 80S\ FPKM}{5'\,UTR\ \ RNA\ FPKM}$$

Scanning efficiency (SE):

$$\frac{5'\,UTR\ SSU\ FPKM}{CDS\ RNA\ FPKM}$$

Initiation rate (IR):

$$\frac{CDS\ 80S\ FPKM}{5'\,UTR\ SSU\ FPKM}$$

Translational efficiency (TE):

$$\frac{CDS\ 80S\ FPKM}{CDS\ RNA\ FPKM}$$

Scanning efficiency and initiation rate are equivalent to translational efficiency:

$$\frac{5'\,UTR\ SSU\ FPKM}{CDS\ RNA\ FPKM} \cdot \frac{CDS\ 80S\ FPKM}{5'\,UTR\ SSU\ FPKM} = \frac{CDS\ 80S\ FPKM}{CDS\ RNA\ FPKM}$$

$$SE\ \cdot\ IR\ =\ TE$$

***Figure S15: Metrics used to investigate interactions of small subunit and 80S footprints.***

#### Supplementary Tables

***Table S1: Small subunit ratio between start and end of over 5' UTR regions.***

| **Transcript Group** | **Median Ratio** | **Count** |
|---|---|---|
| All transcripts | 0.682 | 4969 |
| >=1 ATG uORF | 0.667 | 4738 |
| 0 ATG uORF | 0.951 | 231 |
| The median ratio of small subunit RCP fragments in regions proximal to the start of the transcript or the start codon, for all protein coding transcripts with >= 10 RNA-seq FPKM and >= 220 nt 5' UTRs. | | |

*Table S2: GO analysis summary statistics.*

| GO term | Description | p-value | FDR q-value | Enrichment (N, B, n, b) |
|---|---|---|---|---|
| GO:0044425 | membrane part | $3.64 \times 10^{-14}$ | $2.96 \times 10^{-11}$ | 1.34 (2489,634,995,339) |
| GO:0031224 | intrinsic component of membrane | $2.18 \times 10^{-13}$ | $8.84 \times 10^{-11}$ | 1.38 (2489,544,995,300) |
| GO:0016021 | integral component of membrane | $2.34 \times 10^{-13}$ | $6.33 \times 10^{-11}$ | 1.38 (2489,542,995,299) |
| GO:0016020 | membrane | $9.83 \times 10^{-10}$ | $1.99 \times 10^{-7}$ | 1.26 (2489,713,995,358) |
| GO:0005783 | endoplasmic reticulum | $3.1 \times 10^{-7}$ | $5.03 \times 10^{-5}$ | 1.86 (2489,98,794,58) |
| GO:0005789 | endoplasmic reticulum membrane | $5.48 \times 10^{-6}$ | $7.42 \times 10^{-4}$ | 2.15 (2489,48,773,32) |
| GO:0044432 | endoplasmic reticulum part | $8.13 \times 10^{-6}$ | $9.43 \times 10^{-4}$ | 1.88 (2489,77,773,45) |
| GO:0044444 | cytoplasmic part | $6.37 \times 10^{-5}$ | $6.47 \times 10^{-3}$ | 1.24 (2489,649,718,233) |
| GO:0005793 | endoplasmic reticulum-Golgi intermediate compartment | $5.45 \times 10^{-4}$ | $4.92 \times 10^{-2}$ | 6.96 (2489,6,298,5) |
| Summary statistics of GO analysis for cellular component GO terms based on ranking of genes according to Initiation Rate (IR). P-values computed according to the mHG model. FDR q-value is the correction of the p-value for multiple testing using the Benjamini and Hochberg (1995) method. Enrichment (N, B, n, b) is defined as: N - total number of genes, B - total number of genes associated with a specific GO term, n - number of genes in the top of input list, b - number of genes in the intersection, Enrichment = (b/n) / (B/N). GO analysis was performed with GOrilla [3]. | | | | |

***Table S3: RCP-seq datasets.***

| **Sample** | **Fractions** |
|---|---|
| 64-cell 1 | SSU, LSU |
| 64-cell 2 | SSU, LSU |
| Sphere 1 | SSU, LSU |
| Sphere 2 | SSU, LSU |
| Sphere 3 | SSU, LSU |
| Shield 1 | SSU, LSU<br>f9, f10, f12, f13, f14, f15, f16, f17, f18, f19, f20 |
| Shield 2 | SSU, LSU |
| Shield 3 | SSU, LSU |
| 64-cell-4Ei-0.1 | SSU, LSU |
| 64-cell-4Ei-10 | SSU, LSU |
| Shield 4 (150NT) | SSU, LSU |

Summary of the *D. rerio* RCP-seq libraries, listing the fractions and data used in this project (SSU - small ribosomal subunit, LSU - large ribosomal subunit or 80S) as described also in the SRA uploaded files (PRJEB33323). For sample Shield 1 fractions are available as individual files or as merged SSU and LSU fractions.

*Table S4: Summary of datasets used in this study.*

| Dataset | 1 | 2 | 3 | Publication | Accession |
|---|---|---|---|---|---|
| RCP-seq | 64 cell | Sphere | Shield | - | PRJEB33323 |
| CAGE | 64_cells | Sphere_dome | Shield | Nepal 2013 [4] | SRA055273 |
| Total RNA-seq | 2hpf | 4hpf | 6hpf | Lee 2013 [5] | GSE47558 |
| Ribo-seq | 256cell | Dome | Shield | Chew 2013 [6] | GSE46512 |
| CAGE | Yeast | | | Wery 2016 [2] | GSE69384 |

***Table S5: RCP-seq alignment statistics.***

| **Library** | **Raw reads** | **Reads after adapter trimming** | **Reads mapping to protein coding transcripts** |
|---|---|---|---|
| 64-cell | 277,274,244 | 93,880,163 | 2,485,185 |
| Sphere | 354,199,414 | 146,737,132 | 1,819,780 |
| Shield | 253,786,403 | 165,104,811 | 5,363,177 |
| 64-cell 4Ei-10 0.1µm | 139,688,222 | 85,274,709 | 3,195,345 |
| 64-cell 4Ei-10 10µm | 128,586,905 | 78,315,625 | 2,920,765 |
| Archer 2016 *S. cerevisiae* | 51,381,807 | 34,302,085 | 918,074 |
| Alignment statistics for *D. rerio* RCP-seq small subunit libraries, including *S. cerevisiae* TCP-seq small subunit library for comparison. | | | |

#### Supplementary Notes

**Overview of RCP-seq technique.**

In designing this project, we decided to capture all small subunit complexes and not just those co-existing with translating 80S complexes. This allowed us to study the prevalence of scanning and its efficiency in leading to initiation of translation. We based our choice of crosslinking and sucrose gradients on previously published research in yeast [1,7] and adapted these to zebrafish embryonic samples as described in the methods section.

Successful crosslinking was determined from polysome profiles where the small subunit (40S), monosome peak (80S) and polysome peaks could be observed. Specifically, we could observe a higher small subunit peak in crosslinked vs flash-frozen samples. For separation of the small subunit and 80S complexes, linear gradients from 7 to 30% sucrose gave the best resolution. We observed possible incomplete separation of the fractions in the first sequencing data and therefore chose to include additional fractions before the small subunit peak, in between the small subunit and 80S peaks, and after the 80S peaks (Fig. S14). This allowed us to refine the selected fractions. We chose to sequence the fractions in each peak separately in order to obtain more information about different conformations (three fractions for small subunit complexes and two for 80S complexes). Even though this was not further analysed in this work, we anticipate that with future deeper sequencing, different conformations of the scanning and initiating small ribosomal subunit could be differentiated in those fractions.

In ribosome profiling, typically a size range from only 26 to 34 nt would be selected as "translating" ribosomes. However, this has recently been challenged and it is now believed that different RNA fragment sizes represent different conformations of the ribosomal complex [8]. Except for removing adapter dimers, we did not remove fragments from the library prep by size, as we did not want to select against specific conformations. All sequencing was performed at 75 bp (single-read) with the exception of a single sample which was sequenced at 150 bp. The longer library was constructed to survey all conformations. However, only a few reads were revealed to extend beyond 75bp and based on this it was decided to sequence only 75bp which captures the vast majority of reads and is sufficient to capture the different small subunit conformations.

A limitation in this type of studies is the large presence of rRNA fragments in the resulting sequencing data. For this study, we tested removal of such fragments by targeted rRNA removal through specially designed probes. However, this resulted in incomplete removal of contaminants and removal of many protein-coding transcripts. Therefore, we further tested removal with commercially available kits and opted for the ribo-zero rRNA removal kit, which showed no removal of protein-coding transcripts and an improved removal of rRNA contaminants.

**Effects of 4Ei-10 treatment.**

At the higher concentration of 4Ei-10 (10 μM), approximately half of the embryos were arrested before reaching the shield stage. For those who survived, they suffered severe developmental delays and would eventually disintegrate before 24 hpf. Surprisingly, these severe effects were not reflected in the polysome profiles at Shield stage where there was only a small reduction in polysome peaks (Fig. S6). The monosome peaks remained almost the same, while there was a clear reduction in the more actively translated RNAs as reflected in reduction of the polysome fraction.

Intriguingly, in the absence of threading (4Ei-10 treated samples), we observed an enrichment of shorter fragments over the 5' UTR (RCP-seq data). The length of these corresponded to the length of the 80S translating ribosomes, but lacked the distinct periodicity observed over CDSs in the 80S fraction. These occurred across the whole leader with no specificity to uORFs. A possibility is that these represent 43S PIC recruited to the 5' UTRs through an alternative pathway but lacking most of the canonical factors that are thought to be the cause of longer footprints [1]. These fragments are also detectable in untreated samples, but at a much lower frequency (Fig. S3). If these fragments represent an alternative scanning pathway, it appears to be incapable of compensating for threading as eIF4E inhibition results in severe developmental effects in zebrafish embryos.

We found that the 5’ end peak representative of threading in transcripts to be largely removed after 4Ei-10 treatment. In order to quantify this loss, the ratio of reads at the 5’ peak relative to reads internal to the 5’ UTR was calculated. A limitation in this calculation is the need for a high number of reads from the small subunit in the 5’ UTR. Due to the strong abolishment of scanning and translation in the higher concentration of 4Ei-10 (10μM), most transcripts only have a single read in the leader sequence, making this calculation inaccurate. In the lower concentration treatment (0.1μM), the translational reduction is smaller and more reads are available. However, if the translating length fragments in the small subunit footprints are kept, enough reads are available in both sets of treatments that the 5’ peak loss can be estimated. Taking all small subunit footprint lengths, we find that the peak in 0.1μM-treated samples to be reduced to 63% and in 10μM-treated samples it is further reduced to 50%. For the results shown in the main text and Fig. S7, only the lower 4Ei-10 concentration was used to quantify this loss, as we were able to estimate more accurately the peak loss without the translating length footprints. Regardless of the data included, very similar quantifications of peak loss were obtained.

**Interpretation of footprint densities.**

Footprinting techniques such as ribo-seq, TCP-seq and RCP-seq measure the average number of ribosomes across a given transcript. This measure of ribosome occupancy is commonly used in metrics such as translational efficiency to assess the translational level of transcripts [9]. However it is important to highlight that these techniques do not measure the rate of translation of those ribosomes, and assumptions that an increased ribosomal occupancy leads to increased translation do not always hold [10,11]. For example, accumulations of ribosome occupancy would be expected to occur upstream of transcript features associated with reduced translation rate [12]. The increased ribosome occupancy of such stalled or queued ribosomes could therefore be incorrectly interpreted as an overall increased rate of translation. This means that using these metrics as a measure of protein synthesis should be taken with caution. This may be particularly pertinent to the small subunit complex fractions of RCP-seq datasets, because a number of questions remain regarding the mechanisms of ribosome scanning, and how scanning ribosomes might interact with mRNA features or other ribosomes [13].
